# Myostatin is a major endocrine driver of follicle-stimulating hormone synthesis

**DOI:** 10.1101/2023.08.30.555595

**Authors:** Luisina Ongaro, Xiang Zhou, Ying Wang, Ziyue Zhou, Hailey Schultz, Evan R.S. Buddle, Emilie Brûlé, Yeu-Farn Lin, Gauthier Schang, Roselyne Castonguay, Yewei Liu, Gloria H. Su, Nabil Seidah, Kevin C. Ray, Seth J. Karp, Ulrich Boehm, Se-Jin Lee, Daniel J. Bernard

## Abstract

Myostatin is a paracrine myokine that regulates muscle mass in a variety of species, including humans. Here, we report a functional role for myostatin as an endocrine hormone directly promoting pituitary follicle-stimulating hormone (FSH) synthesis and thereby ovarian function. Previously, this FSH-stimulating role was attributed to other members of the transforming growth factor β family, the activins. The results both challenge activin’s eponymous role in FSH synthesis and establish an endocrine axis between skeletal muscle and the pituitary gland. The data also suggest that efforts to antagonize myostatin to treat muscle wasting disorders may have unintended consequences on fertility.

**One-Sentence Summary:** Hormone synthesis and reproduction depend on crosstalk between skeletal muscle and the pituitary gland.

## Main Text

Follicle-stimulating hormone (FSH) is a dimeric glycoprotein produced by gonadotrope cells of the anterior pituitary gland. Though FSH was recently implicated in post-menopausal bone loss, weight gain, and cognitive decline (*1-3*), its best-known actions are in the gonads (*4*). FSH is a critical regulator of ovarian follicle growth and spermatogenesis. Purified and recombinant forms of the hormone are used for controlled ovarian stimulation in infertile women and to treat hypogonadism in some men (*5, 6*).

FSH is regulated by peptide, protein, and steroid hormones in the hypothalamic-pituitary-gonadal axis (*7*). Gonadotropin-releasing hormone (GnRH) from the brain stimulates the synthesis and secretion of both FSH and the related luteinizing hormone (LH). FSH and LH work in tandem to control gonadal function, including hormone production. Sex steroids, like estradiol and testosterone, provide feedback to the brain and pituitary to primarily inhibit FSH and LH. In response to FSH, the gonads also produce inhibins, TGFβ family ligands that feed back on pituitary gonadotropes to selectively suppress FSH (*7*).

Inhibin A and B were initially purified from ovarian follicular fluid (*8*). The inhibins share an α subunit and are distinguished by their unique β subunits, inhibin βA for inhibin A and inhibin βB for inhibin B. During the purification of the inhibins, three additional proteins were serendipitously discovered based on their selective regulation of FSH secretion from cultured rat pituitary cells. Given their FSH stimulating actions and structural similarity to inhibins, two of these proteins were called activin A and activin AB (*9*). Activin A is a homodimer of inhibin βA subunits whereas activin AB is an inhibin βA-βB heterodimer. Activin B is a homodimer of inhibin βB (encoded by the *Inhbb* gene) subunits, but was not purified from follicular fluid. Activins can stimulate FSH production by binding and signaling through complexes of activin type II (ACVR2A and ACVR2B) and type I receptors (ACVR1B and ACVR1C) to drive transcription of the FSHβ subunit gene (*Fshb*) (*7, 10*, and **Fig. 1A**). FSHβ synthesis is rate-limiting in production of the dimeric hormone. Its α subunit (*Cga*) is not regulated by activins. Inhibins are endogenous antagonists, competing for binding to type II receptors on pituitary gonadotropes (*11, 12*).

**Fig. 1:**
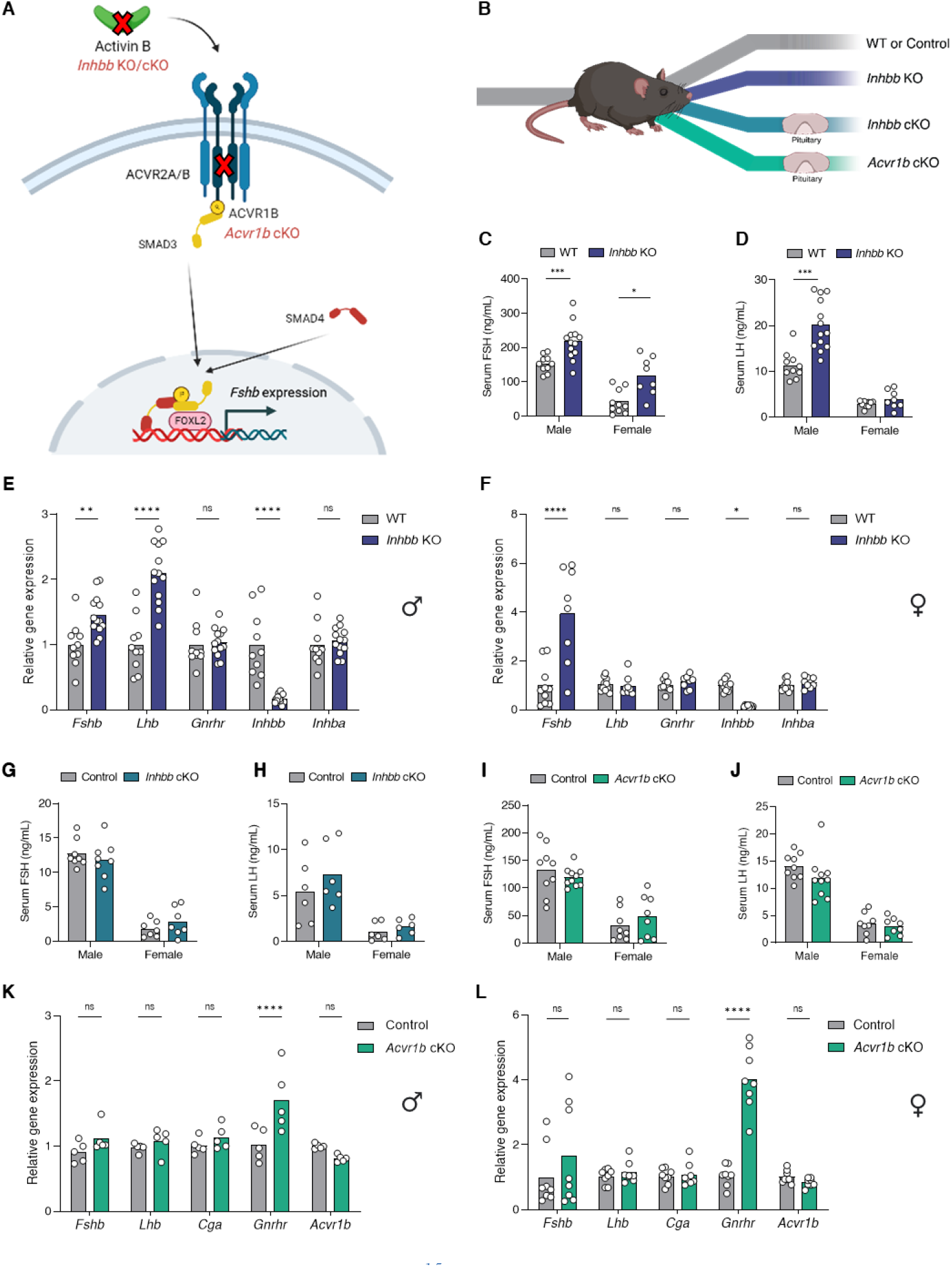
FSH synthesis is activin-independent in mice. **(A)** Schematic representation of the proposed mode of activin B signaling in gonadotropes. The Xs represent the activin B knockout and type I receptor, ACVR1B, knockout models examined in Figure 1. **(B)** Schematic of mouse models used to assess activin signaling in gonadotropes, including wild-type (WT), control (floxed alleles only), *Inhbb* global KO, gonadotrope-specific *Inhbb* cKO, and gonadotrope-specific *Acvr1b* cKO mice. Serum **(C)** FSH and **(D)** LH levels (measured by multiplex ELISA) in wild-type or *Inhbb* KO mice. Pituitary gene expression in **(E)** male and **(F)** female wild-type (WT) or *Inhbb* KO mice. Serum **(G)** FSH and **(H)** LH levels (measured by ELISA) in control or gonadotrope-specific *Inhbb* cKO mice. Serum **(I)** FSH and **(J)** LH levels (multiplex ELISA) in control or gonadotrope-specific *Acvr1b* cKO mice. Pituitary gene expression in **(K)** male and **(L)** female control or *Acvr1b* cKO mice. Animals were sacrificed at 8 to 10 weeks old. Females were sampled at 7 am on estrus morning. Bar heights are group means. Each circle represents an individual mouse. *Rpl19* was used as a housekeeping gene in E, F, K, and L. * *P* < 0.05, ** *P* < 0.01, *** *P* < 0.001, **** *P* < 0.0001. ns, non-significant.

The third protein purified from follicular fluid was named follistatin, given its ability to inhibit FSH secretion (*13*). Unlike the inhibins, follistatins bind and sequester activins, arguably explaining their FSH suppressing activity (*14*). Subsequently, follistatins were shown to neutralize additional TGFβ ligands, in particular myostatin (also known as growth differentiation factor 8 or GDF8) and the related GDF11 (*15*). Notably, myostatin and GDF11 bind activin type II receptors. They can also signal via activin type I receptors, but preferentially use the TGFβ type I receptor, TGFBR1 (*16*).

Though activins are produced in the gonads and are found in circulation, they are largely bound and inactivated by follistatins (*17*). Therefore, it is currently thought that activins produced locally in the pituitary, and particularly activin B by gonadotropes, are the main drivers of FSH synthesis in vivo (*18, 19*). However, this model is based on a relatively small number of observations. First, gene and protein expression studies revealed inhibin βB, but not βA subunit expression in rat gonadotrope cells (*20*). Second, activin B, but not activin A, neutralizing antibodies selectively suppressed FSH synthesis and secretion by cultured rat pituitary cells (*21*). Other data challenge the necessity for activin B in FSH synthesis. For example, the same activin A and B neutralizing antibodies used in rats did not affect FSH production in mouse pituitary cultures (*22*). In contrast, follistatin and a small molecule inhibitor of activin type I receptors, SB431542, blocked FSH production in these cultures (*23*). Moreover, inhibin βB (*Inhbb*) knockout mice, which cannot make activin B, reportedly have elevated rather than the predicted reduction in circulating FSH (*24*). Importantly, however, mice lacking the activin type II receptors in gonadotropes are FSH-deficient (*25*). Considering these observations, we asked whether another TGFβ ligand, in addition to or instead of autocrine/paracrine activin B, drives FSH production in vivo.

### FSH production is activin B-independent in mice

Consistent with an earlier report (*24*), we observed significantly elevated serum FSH levels in female and male *Inhbb* knockout relative to wild-type mice **(Fig. 1A-C)**. Serum LH was also increased in *Inhbb* knockout males, but not females **(Fig. 1D)**. These changes in hormone levels were associated with increased expression of pituitary *Fshb* in both sexes and LHβ (*Lhb*) in males **(Fig. 1E-F)**. As expected, *Inhbb* mRNA levels were reduced in the knockouts. There was no compensatory increase in pituitary inhibin βA (*Inhba*) expression **(Fig. 1E-F)**.

Gonadotrope-derived activin B is thought to be the principal driver of FSH synthesis (*18, 20, 26*). To test this hypothesis and enable conditional (Cre-mediated) deletion of activin B in gonadotropes, we generated mice harboring a floxed (fx) *Inhbb* allele. Serum FSH and LH levels did not differ between control (*Inhbb*^fx/fx^) and gonadotrope-specific knockouts (*Inhbb*^fx/fx^;*Gnrhr*^GRIC/+^; cKO) **(Fig. 1G-H)**. Litter sizes were also normal in *Inhbb* cKO females **(Fig. S1A)**. *Inhbb*, but not *Fshb, Lhb*, or *Inhba* mRNA levels were reduced in pituitaries of cKOs relative to controls **(Fig. S1B)**. In contrast, when we globally recombined the floxed *Inhbb* allele by crossing to *Gdf9*-iCre, the resulting mice exhibited phenotypes observed in the original *Inhbb* knockout strain, including elevated FSH levels and eyelid malformations **(Fig. S1C-D)**. This confirmed that recombination of floxed *Inhbb* generated a null allele. Collectively, these data indicate that activin B of gonadotrope origin is not required for FSH synthesis in mice. Moreover, elevated FSH levels in global *Inhbb* knockouts likely results from the loss of gonadal inhibin B negative feedback.

### FSH secretion is reduced in mice lacking the type I receptor, TGFBR1, in gonadotropes

As mice lacking activin type II receptors in gonadotropes are FSH deficient (25), one or more TGFβ ligands that use these receptors are required for FSH synthesis in vivo. Single nucleus RNA-seq data (*27*) show that the *Inhba* subunit is mainly expressed in stem cells in the murine pituitary (see also **Fig. 4D**). It was, therefore, possible that activin A from these cells maintained FSH production in *Inhbb* knockout mice. To address this possibility, we conditionally ablated the canonical activin type I receptor, ACVR1B (*28*), in gonadotropes **(Fig. 1A-B**; **Fig. S2)**. This would block signaling in these cells by activin A from any source. Remarkably, serum FSH, serum LH, and pituitary *Fshb* and *Lhb* mRNA levels were unaltered in *Acvr1b* cKOs relative to control littermates **(Fig. 1I-L)**. cKOs did, however, exhibit increased GnRH receptor (*Gnrhr*) expression **(Fig. 1K-L)**, which was previously observed in mice lacking *Acvr2b, Tgfbr3*, or *Smad4* in gonadotropes (*25, 29, 30*). In the conditional *Tgfbr3* knockouts, this increase did not appear to alter GnRH sensitivity (*29*).

Paralleling our data in *Acvr1b* cKOs, activin A or B-stimulated FSH secretion was unaltered in pituitaries from mice lacking the other activin type I receptor, *Acvr1c* (*31*). In cultured wild-type pituitaries, however, a small molecule inhibitor of ACVR1B, ACVR1C, and TGFBR1 (SB431542) blocks *Fshb* expression (*23*). Considering all these results, we asked whether there might be a role for the type I receptor, TGFBR1, in FSH synthesis, even though activins are not thought to signal via this receptor (*32, 33*) and murine gonadotropes are not responsive to TGFβ isoforms due to low or absent expression of the TGFβ type II receptor (TGFBR2) **(Fig. S2** and (*34*)). Remarkably, gonadotrope-specific ablation of *Tgfbr1* led to significant decreases in FSH secretion in both males and females **(Fig. 2A-B)**. Serum LH levels were not altered **(Fig. 2C)**. Pituitary *Fshb* expression trended downward in cKO mice, but this was not statistically significant **(Fig. 2D-E)**. *Gnrhr* expression was increased in *Tgfbr1* cKOs **(Fig. 2D-E)**. Consistent with their lower FSH levels, *Tgfbr1* cKO females had fewer ovarian antral follicles and corpora lutea **(Fig. 2F)** and produced smaller litters than controls **(Fig. 2G)**. There was a small, but significant decrease in testis weight in *Tgfbr1* cKO males **(Fig. 2H)**. We did not detect a significant decrease in *Tgfbr1* levels in whole pituitaries of the cKO mice **(Fig. 2D-E)**. This was also the case for *Acvr1b* levels in *Acvr1b* cKO pituitaries **(Fig. 1K-L)**. However, it must be noted that gonadotropes comprise only 3-5% cells in the pituitary gland and these two receptors are expressed in all lineages **(Fig. S2)**. Therefore, *Acvr1b* and *Tgfbr1* expression, and the efficiency of recombination, in these conditional knockouts must be assessed in purified gonadotropes, which we demonstrate later **(Fig. S3D-G)**.

**Fig. 2:**
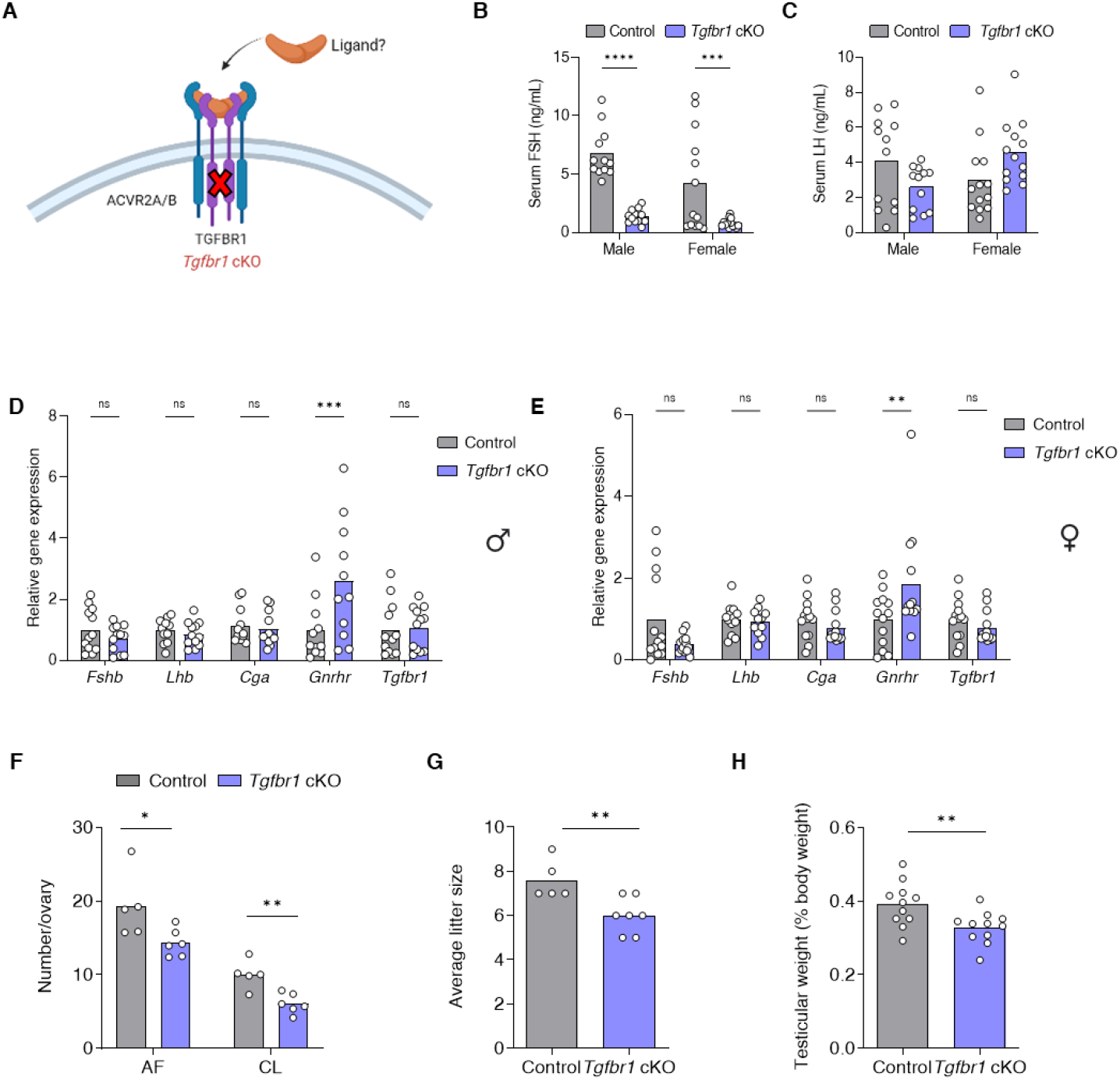
FSH production depends on the type I receptor, TGFBR1, in gonadotropes. **(A)**Schematic representation of the gonadotrope-specific TGFBR1 knockout model used in Figure 2. **(B)** Serum FSH (Luminex assay), **(C)** LH (ELISA), and pituitary gene expression in **(D)** male and **(E)**female control (gray) and gonadotrope-specific *Tgfbr1* cKO (purple) mice. *Rpl19* was used as a housekeeping gene in D and E. **(F)** Antral follicle (AF) and corpora lutea (CL) counts from ovarian sections of 9 to 10-week-old control and *Tgfbr1* cKO females (one ovary per mouse). **(G)** Number of pups per litter in 6-month breeding trials of control and *Tgfbr1* cKO females. **(H)** Testicular weights (normalized to body weight) of control and *Tgfbr1* cKO males. Bar heights are group means. Each circle represents an individual mouse. Females were sampled at 7 am on estrus morning. * *P* < 0.05, ** *P* < 0.01, *** *P* < 0.001, **** *P* < 0.0001. ns, non-significant.

### Mice lacking both ACVR1B and TGFBR1 in gonadotropes are FSH deficient

ACVR2A and ACVR2B show some functional redundancy in murine gonadotropes (*25*). We therefore asked whether the combined ablation of ACVR1B and TGFBR1 might similarly be needed to cause complete FSH deficiency **(Fig. 3A)**. Indeed, FSH was virtually undetectable in serum of gonadotrope-specific *Acvr1b*/*Tgfbr1* knockout mice (double conditional knockouts or dcKOs) **(Fig. 3B)**. As reported in other FSH-deficient models (25, 35), serum LH **(Fig. 3C)** was elevated in female but reduced in male dcKOs. Patterns of pituitary *Fshb* and *Lhb* expression mirrored the hormone levels in circulation **(Fig. 3D-E)**. dcKO females were infertile **(Fig. 3F)**, with reduced ovarian and uterine weights **(Fig. 3G-H, S3A)**. Ovarian folliculogenesis was arrested at the early antral stage in dcKO females **(Fig. 3I)**. Testicular, but not seminal vesicle weights were significantly reduced in male dcKOs **(Fig. S3B-C)**. Testis histology revealed mature spermatozoa in the lumina of the seminiferous tubules of both genotypes **(Fig. 3J)**. The efficacy and specificity of the gene knockouts was confirmed in gonadotropes purified from the animals **(Fig. S3D-G)**. Collectively, these data indicate that FSH is regulated by a TGFβ ligand (or ligands) that signals preferentially via the type I receptor, TGFBR1, with some compensation by ACVR1B.

**Fig. 3:**
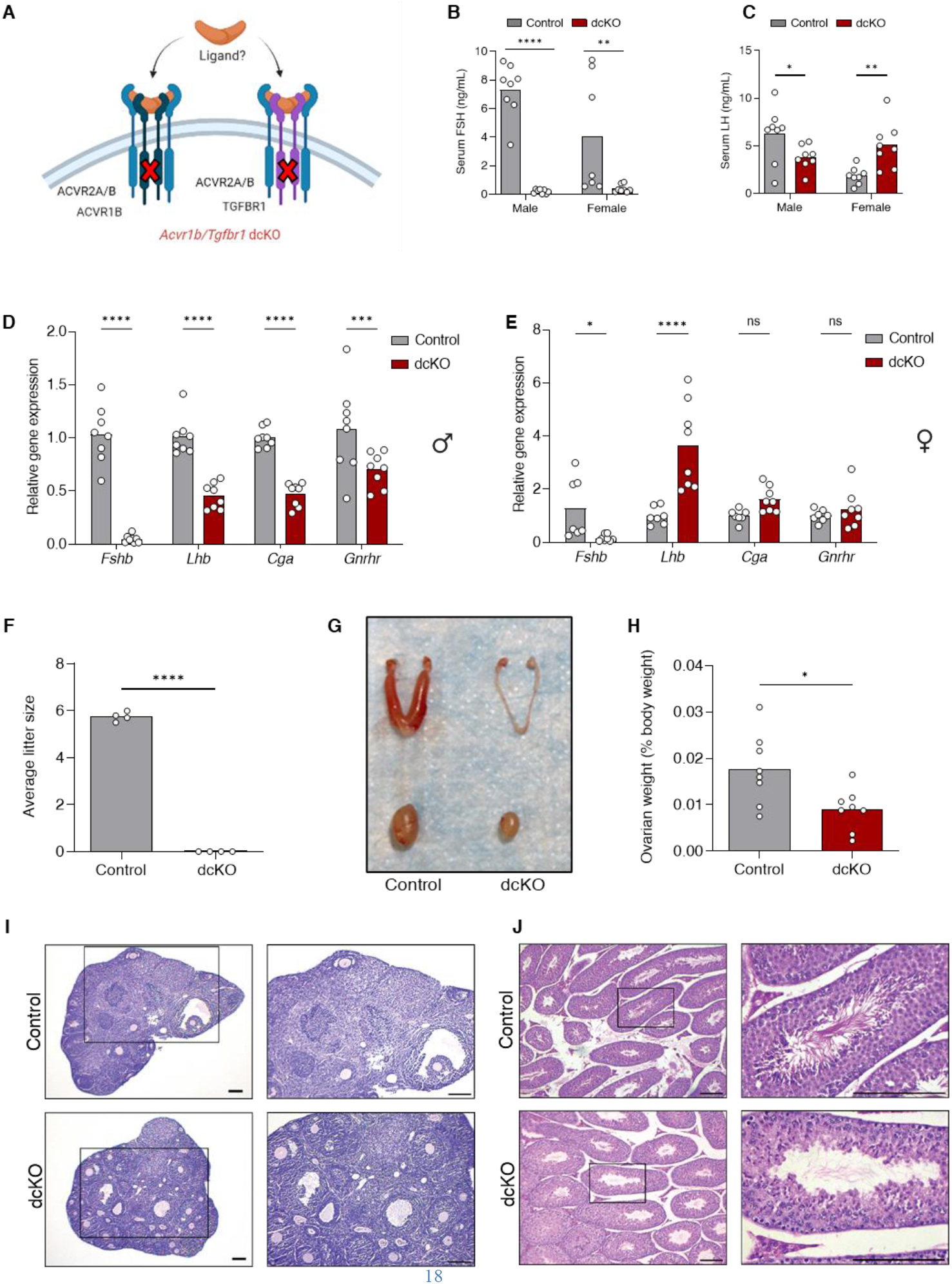
*Acvr1b/Tgfbr1* double cKO animals are FSH-deficient and hypogonadal. **(A)** Schematic representation of the gonadotrope-specific ACVR1B/TGFBR1 knockout model used in Figure 3. Serum **(B)** FSH and **(C)** LH levels (measured by ELISA), and pituitary gene expression in **(D)** male and **(E)** female 9-10-week-old control (gray) and gonadotrope-specific *Acvr1b/Tgfbr1* dcKO (maroon) mice. Control females were randomly cycling. *Rpl19* was used as a housekeeping gene in D and E. **(F)** Number of pups per litter in 6-month breeding trials of control and *Acvr1b/Tgfbr1* dcKO females. **(G)** Representative images of female reproductive tracts (top) and testes (bottom). **(H)** Ovarian weights (normalized to body weight). Bar heights are group means. Each circle represents an individual mouse. * *P* < 0.05, ** *P* < 0.01, *** *P* < 0.001, **** *P* < 0.0001. ns, non-significant. **(I)** Ovarian and **(J)** testicular histology sections stained with H&E. Boxed areas in the images at the left are expanded at the right. Scale bars: 200 µm.

### The TGFβ ligand regulating FSH synthesis is not produced in gonadotropes

Though activin B is regarded as an, if not the, autocrine/paracrine regulator of FSH, the data presented thus far demonstrate that activin B from gonadotropes or elsewhere is dispensable for FSH production in mice. Moreover, whichever TGFβ family member(s) regulate FSH, they must bind to ACVR2A and ACVR2B (*25*), signal via TGFBR1 and ACVR1B, and be antagonized by follistatins. Two non-activin ligands that satisfy these criteria are myostatin and GDF11; however, neither was previously implicated in FSH regulation. Therefore, we first examined the actions of these ligands in the immortalized gonadotrope-like cell line, LβT2. Both myostatin and GDF11 stimulated murine *Fshb* promoter-luciferase (luc) activity **(Fig. 4A)**. These responses were dependent on endogenous ACVR2A, as knocking down the receptor with a previously validated siRNA (*23*) impaired ligand-stimulated reporter activity **(Fig. 4B-C)**. ACVR2A knockdown similarly blocked activin B action in these cells **(Fig. S4A)**. We next validated siRNAs against *Acvr1b* and *Tgfbr1* **(Fig. S4B)**. Knockdown of ACVR1B attenuated activin B **(Fig. S4A)**, but not myostatin or GDF11, induction of *Fshb*-luc in LβT2 cells **(Fig. 4B-C)**. In contrast, TGFBR1 knockdown inhibited myostatin and GDF11, but not activin B, activity **(Fig. 4B-C, S4A)**.

**Fig. 4:**
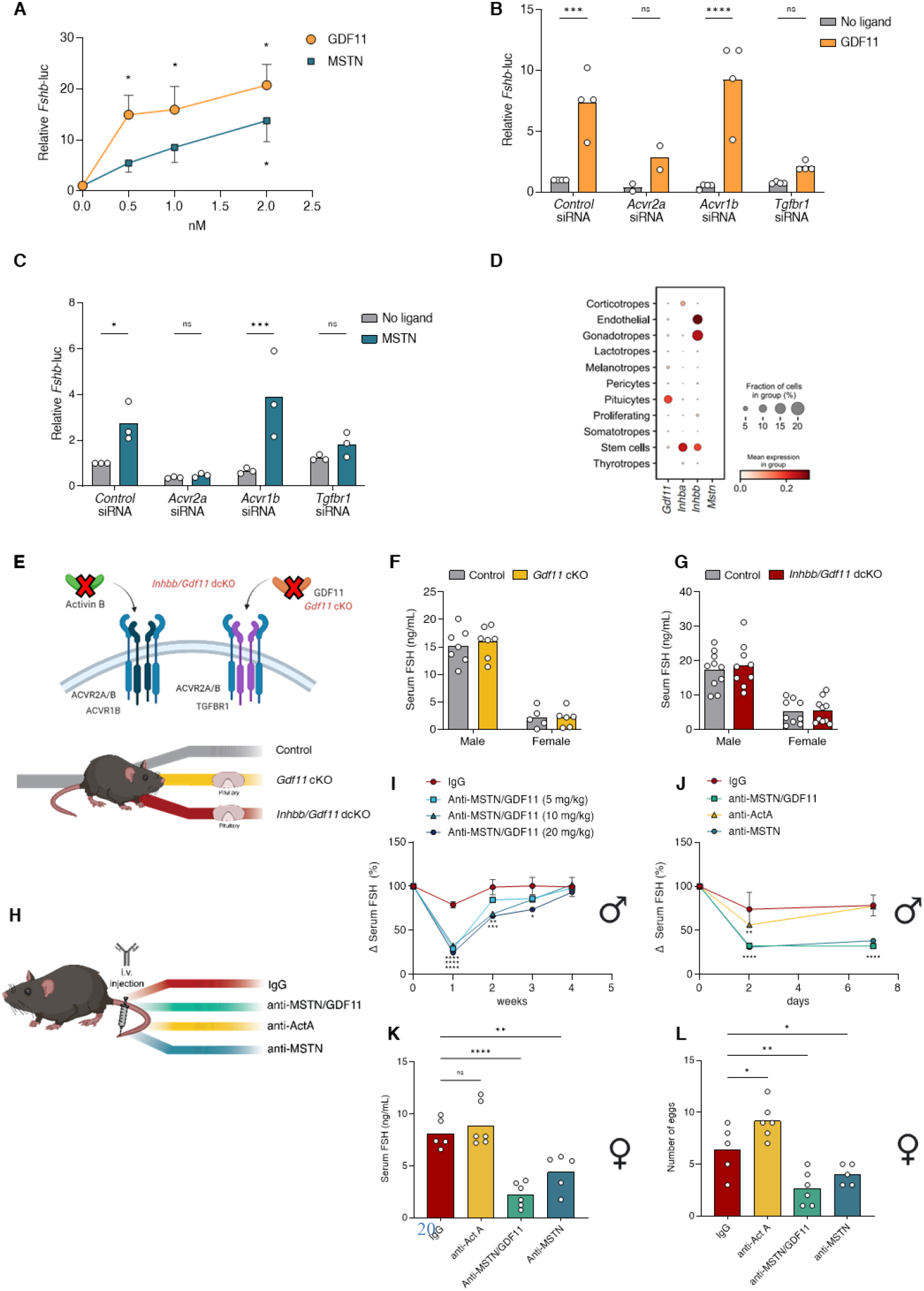
Myostatin (MSTN) and GDF11 stimulate FSH synthesis. **(A)** LβT2 cells were transfected with a murine *Fshb* promoter-luciferase (luc) reporter plasmid. Cells were then treated with media or 0.5-2 nM GDF11 (orange) or MSTN (teal). Data represent mean + or - SEM from 3 independent experiments. **(B-C)** LβT2 cells were transfected with the *Fshb*-luc reporter and 5 nM of control, *Acvr2a, Acvr1b*, or *Tgfbr1* siRNA. Cells were treated with media or **(B)** GDF11 (1 nM) or **(C)** MSTN (2 nM). Bar heights are group means. Each circle represents an independent experiment. **(D)** Dot plots of *Gdf11, Inhba, Inhbb*, and *Mstn* expression in different cell lineages from snRNAseq of adult female and male murine pituitaries. **(E)** Schematic representation of the gonadotrope-specific gonadotrope-specific *Gdf11* and *Inhbb/Gdf11* double knockout mice used in Figure 4F and G. Serum FSH (measured by ELISA) in 9-10-week-old **(F)** control and *Gdf11* cKO or **(G)** control and *Inhbb/Gdf11* dcKO mice. Females were sampled at 7 am on estrus morning. Bar heights are group means. Each circle represents an individual mouse. **(H)** Schematic of intravenous (tail vein) injections of the indicated neutralizing antibodies in adult wild-type mice. **(I)**Percentage change in serum FSH levels 1, 2, 3, and 4 weeks following a single i.v. injection of mouse IgG or 5-20 mg/kg anti-MSTN/GDF11 antibody in adult male wild-type mice. The total amount of injected antibody was balanced to 20 mg/kg using mouse IgG. Data were normalized to FSH levels before injection for each animal. Data represent mean ± SEM (N=3/group). **(J)** Percentage change in serum FSH levels 2 or 7 days following a single i.v. injection of mouse IgG, anti-activin A, anti-MSTN, or anti-MSTN/GDF11 antibodies (10 mg/kg) in adult male wild-type mice. Data were normalized to FSH levels before injection. Data represent mean ± SEM (N=3/group). **(K)** Serum FSH levels (ELISA) and **(L)** number of eggs ovulated by female wild-type mice at natural ovulation following a single i.v. injection with the indicated neutralizing antibodies. Injected females were paired with wild-type males 7 days post-injection and samples collected on the morning of vaginal plugging. Bar heights are group means. Each circle represents an individual mouse. * *P* < 0.05, ** *P* < 0.01, *** *P* < 0.001, **** *P* < 0.0001. ns, non-significant.

Single cell and single nucleus RNA-seq data sets revealed *Gdf11* expression in the murine pituitary gland, including in some gonadotrope cells **(Fig. 4D** and (*27*)). In contrast, myostatin (*Mstn*) was not detected in any pituitary cell type **(Fig. 4D** and (*27*)). We therefore conditionally ablated *Gdf11* alone and in combination with *Inhbb* in gonadotropes in vivo **(Fig. 4E)**. FSH levels were unaltered in *Gdf11* cKO or *Inhbb*/*Gdf11* dcKO mice of either sex **(Fig. 4F-G)**. We confirmed the efficacy and specificity of the gene knockout in purified gonadotropes from dcKO animals **(Fig. S5A-D)**. To assess potential roles for other TGFβ ligands produced by gonadotropes, we conditionally ablated *Furin* in these cells. Furin post-translationally cleaves the prodomains of several TGFβ ligands, a prerequisite for ligand activation (36). FSH production and other reproductive parameters were unaltered in *Furin* cKO males **(Fig. S6A-E)**, despite efficient knockout of the gene **(Fig. S6F-G)**. Females in this line were not assessed. Collectively, these data suggest that activin B, GDF11, or other TGFβ ligands of gonadotrope origin are not required for FSH production in vivo.

### Myostatin is a major regulator of FSH synthesis

Though gonadotrope-derived GDF11 was ruled out in FSH regulation in vivo, this did not preclude a role for the ligand or for myostatin (MSTN) from extra-gonadotrope or extra-pituitary sources. To begin to address this possibility, we treated male wild-type mice i.v. with a MSTN/GDF11 neutralizing antibody (RK-35) (*37*) at 5, 10, or 20 mg/kg **(Fig. 4H)** and measured serum FSH levels 1, 2, 3, and 4 weeks later. After 1 week, FSH was reduced by ~75% at all three doses **(Fig. 4I)**. Levels began to increase in a dose-dependent manner thereafter, with FSH returning to baseline at all three doses by 4 weeks **(Fig. 4I)**. Next, we treated wild-type males with 10 mg/kg of the same MSTN/GDF11 antibody, a MSTN-specific antibody (RK-22), or an activin A neutralizing antibody (SW101) (*37*) **(Fig. 4H, 4J)**. We confirmed the specificity of all three antibodies in vitro **(Fig. S7A-C)**. Both the MSTN/GDF11 and MSTN antibodies reduced FSH in vivo by ~75% within 2 days and the hormone remained at this level at day 7 **(Fig. 4J)**. In contrast, the activin A antibody produced a more modest and shorter-lived response **(Fig. 4J)**. At the conclusion of the experiment (after 7 days), serum FSH and pituitary *Fshb* mRNA levels were equivalently and significantly reduced in the groups treated with the MSTN/GDF11 or MSTN neutralizing antibodies, whereas there was no effect of the activin A antibody **(Fig. S8A-S8B)**. LH levels did not differ significantly between groups **(Fig. S8C)**.

Next, we treated female wild-type mice i.v. with 10 mg/kg of the same antibodies **(Fig. 4H)**. After 7 days, females were paired with untreated wild-type males. On the morning of vaginal plugging (estrus morning; typically within 2-3 days of pairing), serum FSH and pituitary *Fshb* mRNA levels were reduced in females treated with the MSTN/GDF11 or MSTN antibody, though the effect was greater with the former **(Fig. 4K, S8D)**. These lower FSH levels were associated with a reduced number of eggs ovulated **(Fig. 4L)**. The activin A antibody did not significantly affect serum FSH or pituitary *Fshb* but led to a greater number of eggs ovulated **(Fig. 4K-L, S8D)**. The treatments did not significantly affect serum LH or pituitary *Lhb* expression levels at the time of sampling **(Fig. S8E-F)**.

### Myostatin knockout mice have reduced FSH levels and are hypogonadal

To complement the ligand neutralization studies, we next examined reproductive phenotypes in *Mstn* global KO mice (**Fig. 5A**) (*38*). Serum FSH levels and pituitary *Fshb* expression were reduced by about 50% in both male and female (estrus morning) KOs relative to controls **(Fig. 5B-C)**. Female KOs ovulated approximately half the number of eggs as wild type in natural cycles **(Fig. 5D)** and male KOs exhibited a ~40% decrease in testis weights **(Fig. 5E-F)**, even when correcting for their 20% increase in overall body mass (*38*).

**Fig. 5:**
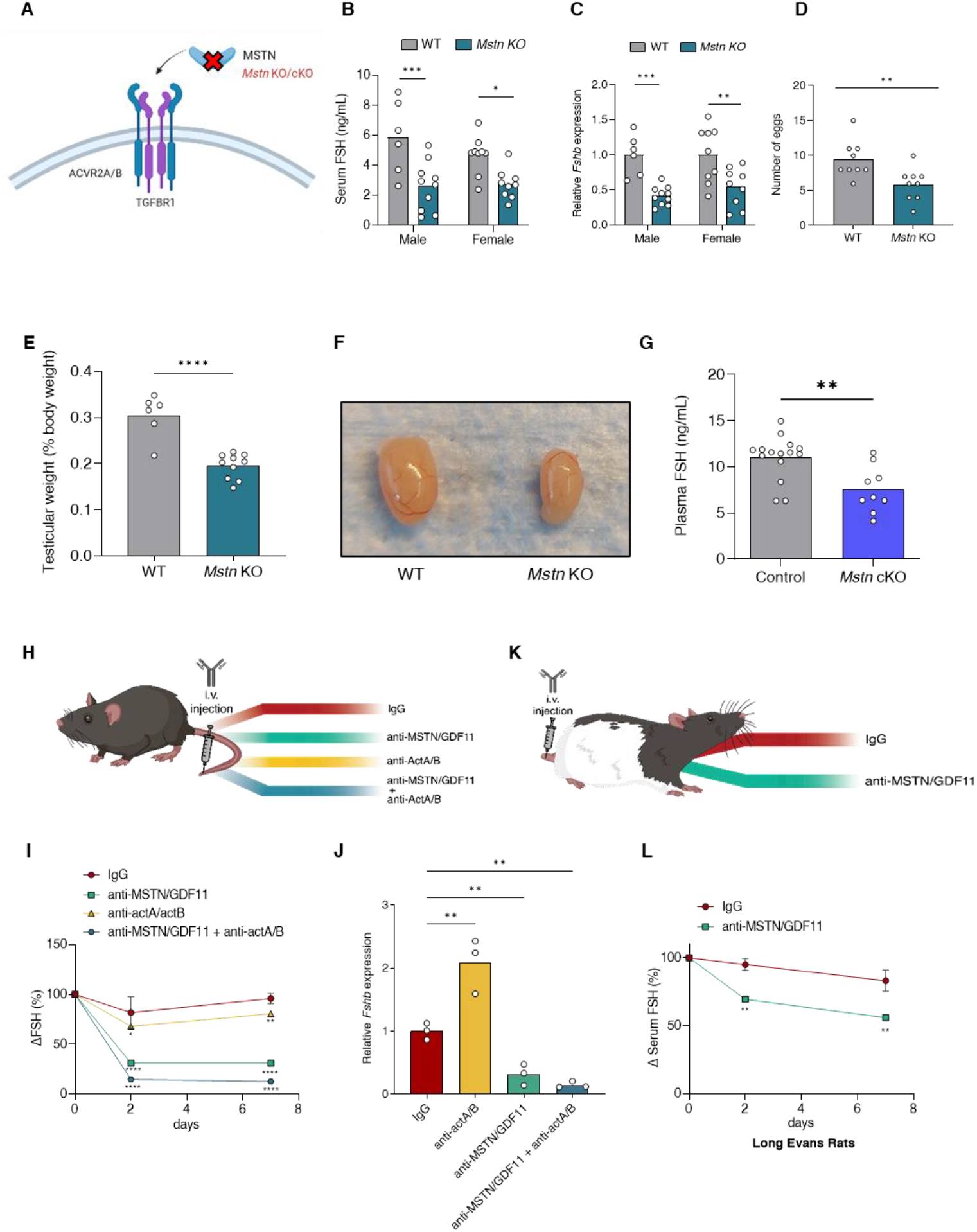
Myostatin is a major driver of FSH synthesis and secretion in mice. **(A)** Schematic representation of *Mstn* knockout mice used in Figure 5B-F. **(B)** Serum FSH (ELISA) and **(C)** pituitary *Fshb* expression in 9-10-week-old control (gray) and *Mstn* KO (teal) mice. Females were sampled at 7 am on estrus morning. *Rpl19* was used as a housekeeping gene in C. **(D)** Number of eggs ovulated on estrus morning in 8-9-week-old control and *Mstn* KO females. **(E)** Testicular weights (normalized to body weight) of 8-week-old control and *Mstn* KO males. In B-E, bar heights are group means. Each circle represents an individual mouse. **(F)** Representative image of testes from a wild-type and a global *Mstn* KO mouse. **(G)** Plasma FSH levels (measured by ELISA) in control or muscle-specific *Mstn* cKO male mice. Plasma was used from archival material in (*12*). Bar heights are group means. Each circle represents an individual mouse. **(H)** Schematic of i.v. injection of neutralizing antibodies in adult wild-type mice. **(I)** Percentage change in serum FSH levels 2 or 7 days following a single i.v. injection of human and mouse IgG, anti-dual activin A/ B (15 mg/kg), and/or anti-MSTN/GDF11 antibodies (10 mg/kg) in adult male wild-type mice. The total amount of injected antibody was balanced across conditions using mouse IgG or human IgG. Data were normalized to FSH levels before injection for each mouse. Data represent mean ± SEM (N=3/group). **(J)** Pituitary *Fshb* expression in adult male mice one week after IgG or antibody injections. Bar heights are group means. Each circle represents an individual mouse. *Rpl19* was used as a housekeeping gene. **(K)** Schematic of i.v. injection of antibodies in adult male Long-Evans rats (200-250 grams). **(L)** Percentage change in serum FSH levels 2 or 7 days following a single i.v. injection of mouse IgG or an anti-MSTN/GDF11 antibody (10 mg/kg) in male rats. Data were normalized to FSH levels before injection for each rat. Data represent mean ± SEM (N=3/group). * *P* < 0.05, ** *P* < 0.01, *** *P* < 0.001, **** *P* < 0.0001. ns, non-significant.

### Muscle-derived myostatin contributes to FSH production

Myostatin is produced predominantly in skeletal muscle (*38*) and circulating levels of the protein are largely derived from this source, as shown in muscle-specific *Mstn* cKO mice (*40*). In residual plasma samples from the mice used in this earlier study, we observed a significant reduction in FSH levels in *Mstn* cKO males relative to controls **(Fig. 5G)**. There was no difference between genotypes in females (data not shown); however, these females were randomly cycling and not sampled on estrus morning, as controlled for in the other models above. Estrus morning is the time of the secondary FSH surge, which depends on the activity of TGFβ ligands (*29*).

### Myostatin is the most critical of four TGFβ ligands driving FSH synthesis in mice

In contrast to *Acvr2a*/*Acvr2b* dcKO (25) or *Acvr1b*/*Tgfbr1* dcKO mice **(Fig. 3B)**, global and muscle-specific *Mstn* KO mice continue to produce some FSH. This suggested that one or more TGFβ ligands may compensate in myostatin’s absence. To assess GDF11’s role, we treated *Mstn* global KOs with the MSTN/GDF11 neutralizing antibody. This further reduced, but did not eliminate, serum FSH and pituitary *Fshb* mRNA levels **(Fig. S9A-C)**, as well as the number of eggs ovulated in females **(Fig. S9D)**. It is important to note that GDF11 levels in circulation do not increase in *Mstn* global KOs (*39*), so this likely does not reflect an insufficiency of the antibody. We therefore examined potential roles for the activins, even though the addition of an activin A neutralizing antibody to the MSTN/GDF11 antibody did not further reduce FSH levels or *Fshb* expression in wild-type male mice **(Fig. S10A-B)** and activin B (*Inhbb*) knockout mice have elevated FSH **(Fig. 1C)**. An antibody with neutralizing activity against both activin A and B (REGN16430, validated in **Fig. S7D)**, nearly eliminated (~90% reduction) serum FSH and pituitary *Fshb* mRNA when used in combination with the MSTN/GDF11 antibody in wild-type males **(Fig. 5H-J)**. Thus, in the absence of myostatin and GDF11, activin A and/or B is/are sufficient to maintain a small level of FSH production in mice.

### Myostatin and GDF11 regulate FSH in rats

The current concept that autocrine/paracrine activin B is the major driver of FSH synthesis derives principally from studies performed in rats (*18, 19*). It was therefore possible that myostatin’s role in mice might be species-specific. Indeed, myostatin levels far exceed those of GDF11 in mouse blood, whereas the two circulate at comparable levels in rats and humans (*41*). In all these species, myostatin and GDF11 circulate in the ng/mL range whereas activin A levels are at least an order of magnitude lower. Equivalent data are not yet available for activin B to our knowledge. We treated adult male rats with 10 mg/kg of the MSTN/GDF11 antibody. As in mice, we observed a significant decrease in FSH levels both 2 and 7 days following the injection **(Fig. 5K-L)**. Unfortunately, we did not have sufficient MSTN/GDF11 antibody for dose-response studies or sufficient MSTN antibody for any studies in rats.

## Discussion

Here, we report the discovery that myostatin is the primary TGFβ family ligand driving FSH synthesis in mice. As myostatin is not produced in the pituitary gland, our results challenge two elements of current dogma. First, an endocrine rather than autocrine TGFβ ligand stimulates FSH. Second, this ligand is not activin B of gonadotrope origin or, for that matter, any activin subtype. It is notable that activins were so named because of their presumed roles as important inducers of FSH synthesis and secretion (*9, 42*). Our data demonstrate that activin A or B, regardless of cellular/tissue origin, only minimally contribute to FSH production or secretion in vivo, and this is evident only when myostatin and/or GDF11 actions are blocked. FSH is greatly reduced in myostatin knockout mice and there is no compensatory increase in circulating GDF11 in these animals (*39*). However, in mice in which GDF11 is knocked into the myostatin locus, circulating levels of GDF11 reach those of myostatin (*39*). In plasma from these animals, we observed a complete rescue of FSH production **(Fig. S10C)**. Thus, myostatin’s dominant role in stimulating FSH is most likely explained by its high levels in circulation, dwarfing those of GDF11 and activins in mice (*41*).

In this regard, it is critical to ask whether myostatin is similarly important in other species. Myostatin’s best known function is in the regulation of muscle mass, and this is the case in a variety of species. Some cattle breeds with so-called double-muscling phenotypes have inactivating mutations in myostatin (*43*). Interestingly, they also exhibit fertility problems (*44*). It is interesting to speculate that these may stem from deficiencies in FSH. Though, it should be noted that myostatin may also have direct actions in uterus, placenta, and ovary (*45*). Whereas myostatin plays a primary role in regulating skeletal muscle in mice and cattle, both myostatin and activin A are critical in primates (*41*). Therefore, in other species, it may be the combinatorial actions of activin-class ligands (i.e., myostatin, GDF11, activin A, activin B, and activin AB) that promote FSH synthesis. Indeed, this is even the case, to some extent, in mice, where neutralization of all these ligands is required for the full suppression of FSH. In rats, we observed a 50% reduction of FSH in response to the MSTN/GDF11 neutralizing antibody, suggesting an additional or compensatory role for other TGFβ ligands. However, as noted, we did not have sufficient antibody to perform dose response experiments in rats. Nevertheless, our data demonstrate that myostatin and/or GDF11 regulate FSH production in both mice and rats. Though we are keen to assess roles for these proteins in humans, we have not yet been able to acquire samples from individuals treated with comparable neutralizing antibodies.

As myostatin is predominantly produced in skeletal muscle, our data suggest a novel endocrine axis in the control of reproduction. That the muscle communicates with pituitary gonadotropes is most clearly demonstrated by the reduced FSH levels in muscle-specific myostatin knockout males. However, it is important to acknowledge that the extent of the FSH loss in these mice was somewhat less than observed in *Mstn* global knockouts, suggesting a potential role for myostatin from additional sources. In this regard, myostatin is expressed in the brain (*46*) and we cannot yet rule in or out a neuroendocrine mode of regulation. Indeed, an FSH releasing factor from the hypothalamus has long been sought but not identified (*47*).

Assuming that muscle-derived myostatin is the primary driver of FSH synthesis, it begs the question of why? We lack a definitive answer. However, as sufficient musculature is required both for reproductive behavior and maintenance of pregnancy, one could speculate that myostatin provides a peripheral signal to the pituitary indicating a physiological level of preparedness. This is analogous to adipose tissue communicating via leptin to the central nervous system to regulate reproductive physiology (*48*). As a test of this idea, it will be important to examine the relationships between muscle mass, circulating myostatin, FSH levels, and reproductive health in humans and other animals. Such analyses should be extended to GDF11 and/or activin A, depending on the species-specific complement of activin-class ligands in circulation coming from skeletal muscle and perhaps other sources.

Finally, our findings may have clinical significance. First, measurements of myostatin, GDF11, and activins, in addition to the inhibins, in circulation should now be considered in assessment of unexplained FSH dysregulation. Second, there are several drug development efforts aimed at inhibiting myostatin activity as a means to treat muscle wasting disorders (*49*). These include myostatin neutralizing antibodies and ligand traps. In the latter case, the ectodomains of ACVR2A and ACVR2B fused to Fc have been developed, tested, and approved for several indications (50). Both of these ligand traps inhibit FSH levels in post-menopausal women (*50, 51*). Though this has been attributed to activin antagonism, our data suggest inhibition of myostatin as an alternative explanation. Moving forward, inhibitory effects of any myostatin antagonist on FSH and fertility should be considered, if not anticipated.

## Acknowledgments

The authors would like to thank the following investigators for providing the indicated reagents: Dr. Teresa Woodruff (rat ACVR1B-HA plasmid, Michigan State University); Dr. Ying Zhang (TGFBR1-Flag plasmid, University of California-Davis); Dr. Thomas Thompson (HEK-CAGA-luc cells, University of Cincinnati); Dr. Terry Hébert (HEK 293 cells; McGill University), Dr. Pamela Mellon (LβT2 cells; University of California, San Diego). We also thank Julie Lord (Flow Cytometry Core, Montreal Clinical Research Institute) for her help with fluorescence-activated cell sorting.

## Author contributions

Conceptualization: DJB, LO, SJL

Methodology: DJB, LO, XZ, YW, ZZ, HS, EB, YFL, GS

Investigation: LO, XZ, YW, ZZ, EB, YFL,

GS Software Programming: HS, ERSB

Resources: RC, GS, NS, KR, SK, UB, SJL

Funding acquisition: DJB

Writing – original draft: LO, HS, DJB

Writing – review & editing: DJB, LO

## Data and materials availability

All data are available in the main text or the supplementary materials. Antibodies against MSTN/GDF11 (RK35), MSTN (RK22), and activin A (SW101) were generously provided by Merck & Co., Inc., Rahway, NJ, USA. A human IgG4 antibody (REGN1945) and an antibody against activin A and B (REGN16430) were generously provided by Regeneron Pharmaceuticals Inc, Tarrytown, NY, USA).

## Supplementary Materials

Materials and Methods

Figs. S1 to S10

Tables S1 and S2

References (*52–62*)

