## Supplementary files for "Myostatin is a major endocrine driver of follicle-stimulating hormone synthesis"

### Materials and Methods

#### Animals

Global *Inhbb* and *Mstn* knockout (KO) mice were described in (*24, 38*). Heterozygous animals were crossed inter-se to produce wild-type or KO mice. *Gnrhr*-IRES-Cre (GRIC) mice (*52*) were used to generate gonadotrope-specific (conditional) knockouts (cKO), with the following floxed (fx) alleles: *Acvr1b* (*53*), *Tgfbr1 (54)*, *Gdf11* (*55*), *Furin* (*56*), and *Inhbb* (*Inhbb*$^{tm1c(EUCOMM)Hmgu}$; MGI:7442209; see below).

Mice carrying *Inhbb* floxed alleles were produced from engineered embryonic stem (ES) cells (JM8.N4; *Inhbbt*m1a(EUCOMM)Hmgu; MGI: 4436214; European Mouse Mutant Cell Repository). ES cell clones were injected into Albino C57BL6/J mice at the Vanderbilt-Ingram Cancer Center (CA68485), Diabetes Research and Training Center (DK20593), and the Vanderbilt Brain Institute in accordance with institutional and federal guidelines. After backcrossing chimeric mice and screening their progeny for germline transmission, mice positive for the modified allele were crossed with the Flp delete mice (B6.Cg-Tg(ACTFLPe)9205Dym/J; RRID:IMSR_JAX:005703) to excise the floxed Neo cassette. After segregating the *Inhbb*$^{fx}$ and Flp alleles, *Inhbb*$^{fx/+}$ males and females were intercrossed to generate *Inhbb*$^{fx/fx}$ homozygotes. These animals were fertile and phenotypically normal. To generate a globally recombined *Inhbb* allele (*Inhbb*$^{\Delta}$), floxed *Inhbb* males were crossed to i*Gdf9*-Cre females (*57*).

Genotyping and assessment of genomic recombination of each floxed, recombined, or wild-type allele were conducted as previously described; see references and primers in **Table S1**.

Seven-week-old (~200-250 g) Long Evans male rats (stain code: 006) and 7-week-old wild-type male and female mice (C57BL/6, strain number: 027) were purchased from Charles River Labs (Senneville, Quebec, Canada). Animals were acclimatized for one week before i.v. injections.

Animals were housed on a 12:12 light/dark cycle (lights on at 7:00) and provided with food and water ad libitum. All animal experiments were performed in accordance with institutional and federal guidelines and were approved by the McGill University and Goodman Cancer Centre Facility Animal Care Committee DOW-A (Protocol 5204).

#### Gonadotrope purification

Gonadotropes were labelled by crossing floxed mice with a *Gt(ROSA26)*$^{ACTB\text{-}tdTomato\text{-}EGFP}$ mice (mTmG, strain number: 007676 from Jackson Laboratories) and sorted by fluorescence-activated cell sorting (FACS) as described previously (*25*). FACS of gonadotropes was performed at the Flow Cytometry Core at the Montreal Clinical Research Institute. For pituitary cell dispersion, 3 pituitary glands per genotype were digested and filtered (0.44 µM). Cell yields were ~$2.0 \times 10^4$ EGFP-positive (gonadotropes) and ~$4.4 \times 10^5$ tdTomato-positive (non-gonadotropes) cells per sample. Following sorting, cells were lysed, and RNA was extracted by column according to the manufacturer's protocol (Geneaid, RB300, New Taipei City, Taiwan). For DNA isolation, after RNA extraction was completed, 50 µl of 8 mM NaOH was added to the columns and incubated at 55C for 15 minutes. Following centrifugation, PCR was performed to assess recombination of the floxed alleles. The primers used were previously described, see references and sequences in **Table S1**.

### Blood collection and hormone analysis

Blood collected from 8- to 10-week-old males or 9- to 11-week-old females by cardiac puncture was allowed to coagulate at room temperature for 30 min and spun down for 15 min at 800 × g. Serum was collected and stored at -20°C. Serum FSH was measured using multiplex ELISA at the Ligand Assay and Analysis Core of the Center for Research in Reproduction at the University of Virginia (Charlottesville, VA, USA), Luminex (Millipore; MPTMAG-49K), or an in-house ELISA (*58*). Serum LH was assessed by multiplex assay at the Ligand Assay and Analysis Core of the Center for Research in Reproduction at the University of Virginia (Charlottesville, VA, USA) or by an in-house ELISA (*59*).

Serum or plasma FSH and LH levels were also measured by in-house ELISAs in muscle-specific *Mstn* cKO mice (*40*) and animals carrying a modify *Mstn* allele in which the mature MSTN C-terminal peptide was replaced with the corresponding region of GDF11 (*Gdf11*/+ or *Gdf11*/*Gdf11*) (*39*).

### Tissue collection and histology

At sacrifice, different organs were dissected, including: pituitary glands, ovaries, uteri, testes, and seminal vesicles. Organs were weighed on an analytical balance or snap frozen in liquid nitrogen for RNA extraction. For histology, one testis per mouse was immersed in Bouin's Fixative Solution (1120-16, Ricca Chemical Company, Arlington, TX, USA) overnight at room temperature, followed by 24 h incubation in 100% ethanol. Testes were stored in 70% ethanol. Two fixed testes from each genotype were paraffin-embedded, sectioned, and stained with hematoxylin/eosin (H&E) at the McGill Centre for Bone and Periodontal Research. Ovaries were fixed in 10% neutral buffered formalin (NBF; HT501128, MilliporeSigma, Burlington, MA, USA) overnight at room temperature, and then stored in 70% ethanol. Fixed ovaries were paraffin-embedded, sectioned, and stained with H&E. All H&E images were acquired with a Leica Microsystems DFC310 FC1.4-megapixel digital color camera on a Leica Microsystems DM1000 light-emitting diode microscope. Follicle counting was performed on fully cut-through ovaries.

### RNA extraction and reverse transcription quantitative PCR (RT-qPCR)

RNA was extracted from pituitary glands using TRIzol Reagent (15596018; Invitrogen, Waltham, MA, USA) following the manufacturer's protocol. For each sample, 200 ng of total RNA were reverse transcribed using random hexamers (C1181, Promega) and M-MLV reverse transcriptase (M1701, Promega, Madison, WI, USA). The resulting cDNA was used for qPCR analysis using EvaGreen (ABMMmix, Diamed, Missisauga, ON, CAN) and primers previously described (see references and sequences in **Table S2**) on a Corbett Rotorgene 600 instrument (Corbett Life Science, Sydney, NSW, AUS). mRNA levels were determined using the $2^{-\Delta\Delta Ct}$ method. Gene expression was normalized to ribosomal protein L19 (*Rpl19*). All primers were validated for efficiency and specificity.

### Breeding trials

At 9 weeks of age, females were paired with wild-type, age-matched C57BL/6 males for 3 to 6 months. Breeding cages were monitored daily to record the number of pups produced. Pups were euthanized 2 weeks after birth.

### Natural ovulation

Randomly cycling, 8- to 9-week-old females were paired with wild-type C57BL/6 males. In experiments involving i.v. antibody treatments, females were paired with males 1 week post-injection. On the morning of vaginal plugging, females were euthanized, and cumulus-oocyte complexes (COCs) were harvested from the ampullae of both oviducts as described in (*12*). COCs were digested with 0.5 mg/mL hyaluronidase (H3884, Millipore Sigma) for 3-5 min at room temperature until dissociated. The number of oocytes was counted using an inverted microscope.

**Single cell analyses**

The murine snRNA-seq dataset and collection details were previously published (*27*). The dataset was downloaded from the GEO data repository (GSE151961). We applied standard preprocessing methods (cell quality filtering, doublet filtering, dimensionality reduction, and scaling) and subsequently integrated and analyzed the datasets using Scanpy 1.9.1 (*60*, *61*). Cell types were assigned as previously published, and Leiden clustering was performed using a resolution of 0.6 (*27*).

**Luciferase reporter assays**

LβT2 cells (provided by Dr. Pamela Mellon, University of California, San Diego, CA, USA) were seeded in 48-well plates at a density of 150,000 cells/well. Cells were cultured 37°C/5% $CO_2$ in DMEM (Dulbecco's Modified Eagle Medium, 319-005-CL, Wisent Inc., Saint-Jean-Baptiste, QC, CAN) with 10% FBS (fetal bovine serum, 098150, Wisent Inc). The following day, cells were transfected overnight with 225 ng/well of the murine −1990/+1 *Fshb* promoter-luciferase reporter *(62)* with or without 5 nM control siGENOME non-targeting (D-001210-05-05), *Acvr2a* (D-040676-02-0005), *Acvr1b* siGENOME (D-043507-03-0005), or *Tgfbr1* siGENOME (D-040617-02-0005) siRNAs (Dharmacon, Lafayette, CO, USA) using Lipofectamine 2000 (L11668019) or Lipofectamine 3000 (L3000015) following the manufacturer's instructions (ThermoFisher Scientific, Waltham, MA, USA). After transfection, cells were incubated overnight in serum-free DMEM. On the following day, cells were treated with activin A (388-AC-050, R&D Systems, Minneapolis, MN, USA), activin B (659-AB, R&D Systems), myostatin (MSTN, 788-G8, R&D Systems) or growth differentiation factor 11 (GDF11, 1958-GD-010, R&D Systems) for 24 h (see final concentrations in figure legends). Cells were then washed with PBS and lysed in 1× passive lysis buffer (PLB, 50 µl/well).

HEK293-$CAGA_{12}$-luciferase reporter cells (CAGA-luc, provided by Dr. Thomas Thompson, University of Cincinnati, OH, USA) were seeded at 50,000 cells/well. Cells were cultured at 37°C/5% $CO_2$ in DMEM with 10% FBS and 100 µg/mL G418 (450-130-QL, Wisent Inc). The next day, activins, MSTN, or GDF11 were pre-incubated for 15 minutes at 37C with murine IgG (Sigma Aldrich I5381), human IgG (REGN 1945, Regeneron Pharmaceuticals Inc, Tarrytown, NY, USA), anti-MSTN (RK22, Merck & Co., Inc., Rahway, NJ, USA), anti-MSTN/GDF11 (RK35, Merck & Co., Inc., Rahway, NJ, USA), anti-activin A (SW101, Merck & Co., Inc., Rahway, NJ, USA), or anti-dual activin A/activin B antibody (REGN 16430, Regeneron). After the incubation, CAGA-luc cells were treated with each complex for 6 hours. Cells were then washed with PBS and lysed in 1× PLB (50 µL/well).

Twenty µL of cell lysis supernatant were combined with 100 µL of assay buffer (15 nM potassium phosphate (pH 7.8), 25 nM glycylglycine, 15 mM magnesium sulfate, 4 nM EDTA, 2 nM adenosine 5′-triphosphate, 1 nM dithiothreitol, and 0.04 mM d-luciferin), and luciferase activity was measured on an Orion II microplate luminometer (Berthold Detection Systems, Oak Ridge, TN, USA). All conditions were performed in technical duplicates or triplicates and the experiments were repeated two to three times.

**siRNA knockdown validation by immunoblotting**

Rat ACVR1B-HA expression plasmid (provided by Dr. Teresa Woodruff, Michigan State University, East Lansing, MI USA) was modified to change 3 bp to those observed in murine *Acvr1b*, providing a perfect match for murine *Acvr1b* siRNA. Here, we used a modified QuikChange protocol with PfuTurbo DNA Polymerase (Agilent Technologies, 200250, Santa Clara, CA, USA) and the following primers (Fw: 5'-CGTAGCTTCTGGTCACATACAACCTTTCGCATTTCCTCAAT-3'/ Rv: 5-ATTGAGGAAATGCGAAAGGTTGTATGTGACCAGAAGCTACG-3'). A rat ALK5-HA expression plasmid was generated in-house from a ALK5-FLAG plasmid (provided by Dr. Ying Zhang, University of California, San Francisco, California, USA) using the following primers (Fw: 5'-AAAAAAGCTTACCATGGAGGCGGCGTCGGCT-3'; Rv: 5'-TTTTATCGATTCCCATTTTGATGCCTTCCTGTTGG-3' and ligation into a C-terminal 3xHA-expressing pcDNA3.0 backbone. The murine *Tgfbr1* siRNA was a perfect match for the corresponding rat *Tgfrb1* sequence.

To assess siRNA knockdown efficiency, HEK293 cells were seeded in a 6-well plate (80,0000 cells/well). Cells were cultured at 37C, 5% $CO_2$ in DMEM with 10% FBS. The following day, cells were transfected with 800 ng per well of either pcDNA3.0, ACVR1B-HA, or TGFBR1-HA expression vector in combination the above indicated siRNAs using 3 µl/well Lipofectamine 3000. Two days post-transfection, cells were lysed in radioimmunoprecipitation assay (RIPA) buffer containing protease inhibitors (cOmplete™ Mini, EDTA-free Protease Inhibitor Cocktail, Roche Holding AG, 04693116001, Basel, Switzerland). Protein concentrations were measured using a Pierce BCA protein assay kit (23227, Thermo Fisher Scientific, Waltham, MA, USA)

Cell lysates were heated in Laemmli buffer (250 mM Tris pH 6.8, 10% SDS, 50% glycerol, 0.2% bromophenol blue, and 10% β-mercaptoethanol) at 70°C for 10 min and resolved by sodium dodecyl sulfate-polyacrylamide gel electrophoresis (SDS-PAGE) on 8% gels prepared using a 30% (w/w) acrylamide/bis-acrylamide (29:1) solution in running buffer (25 mM Tris, 250 mM glycine, 0.1% SDS, pH 8.3). Proteins were transferred to Protran nitrocellulose membranes (GE 10600001, MilliporeSigma) in Towbin buffer (25 mM Tris, 192 mM glycine, pH 8.3, 20% methanol), blocked with 5% skim milk (w/v) in Tris-buffered saline [TBS; 150 mM NaCl, 10 mM Tris (pH 8.0)] containing 0.05% (v/v) Tween 20 (TBST) and incubated overnight at 4°C with agitation with an anti-mouse anti-HA antibody (1:40,000; H9658; RRID:AB_260092, Sigma-Aldrich) or an anti-mouse anti-β-actin (1:40,000; A5441; RRID:AB_476744, Millipore Sigma) diluted in blocking buffer. The following day, membranes were washed in TBST and incubated in horseradish peroxidase-conjugated anti-mouse (1:5,000 goat anti-mouse, 1706516; RRID:AB_11125547, Bio-Rad Laboratories, Hercules, CA, USA) in blocking buffer for 1 h at room temperature with agitation. Membranes were then washed in TBST, incubated in enhanced chemiluminescence substrate (NEL105001, PerkinElmer, Waltham, MA, USA), and bands visualized with an Amersham Imager 600 (GE Healthcare, Chicago, IL, USA).

**In vivo neutralization of activins and/or growth differentiation factors in mice and rats**

For systemic neutralization of activin A, activin B, myostatin (MSTN), and/or GDF11 activities, adult male rats (200 to 250 grams) and adult female or male mice (25 to 30 grams) received a single tail vein injection of 10 mg/kg of murine IgG (Sigma Aldrich, I5381), anti-MSTN (RK-22; Merck & Co., Inc., Rahway, NJ, USA), anti-MSTN/GDF11 (RK-35; Merck & Co., Inc., Rahway, NJ, USA), anti-activin A (SW-101; Merck & Co., Inc., Rahway, NJ, USA) and/or a human IgG4 control antibody (15 mg/kg, REGN1945, Regeneron) or an activin dual-anti activin

A/B antibody (15 mg/kg, REGN16430; Regeneron). Blood (200 µL) was collected from the submandibular venipuncture in mice or the saphenous venipuncture in rats immediately before and 2- and 7-days post-injection. One-week post-injection, male rats or mice were sacrificed for tissue and blood collection. Female mice were paired with wild-type mice to assess natural ovulation, collect organs, and blood (serum). For the RK-35 dose response, adult male wild-type mice were injected via the tail vein with 5, 10, or 20 mg/kg of the antibody. Submandibular blood was collected prior to injection and then weekly for 4 weeks.

**Statistical Analyses**

Depending on the experimental design, data were analyzed by t-test, one-way ANOVA, or two-way ANOVA followed by Holm-Sidak's multiple comparisons test. Statistical analyses were performed using Prism 9.5, GraphPad software. $P < 0.05$ was considered statistically significant.

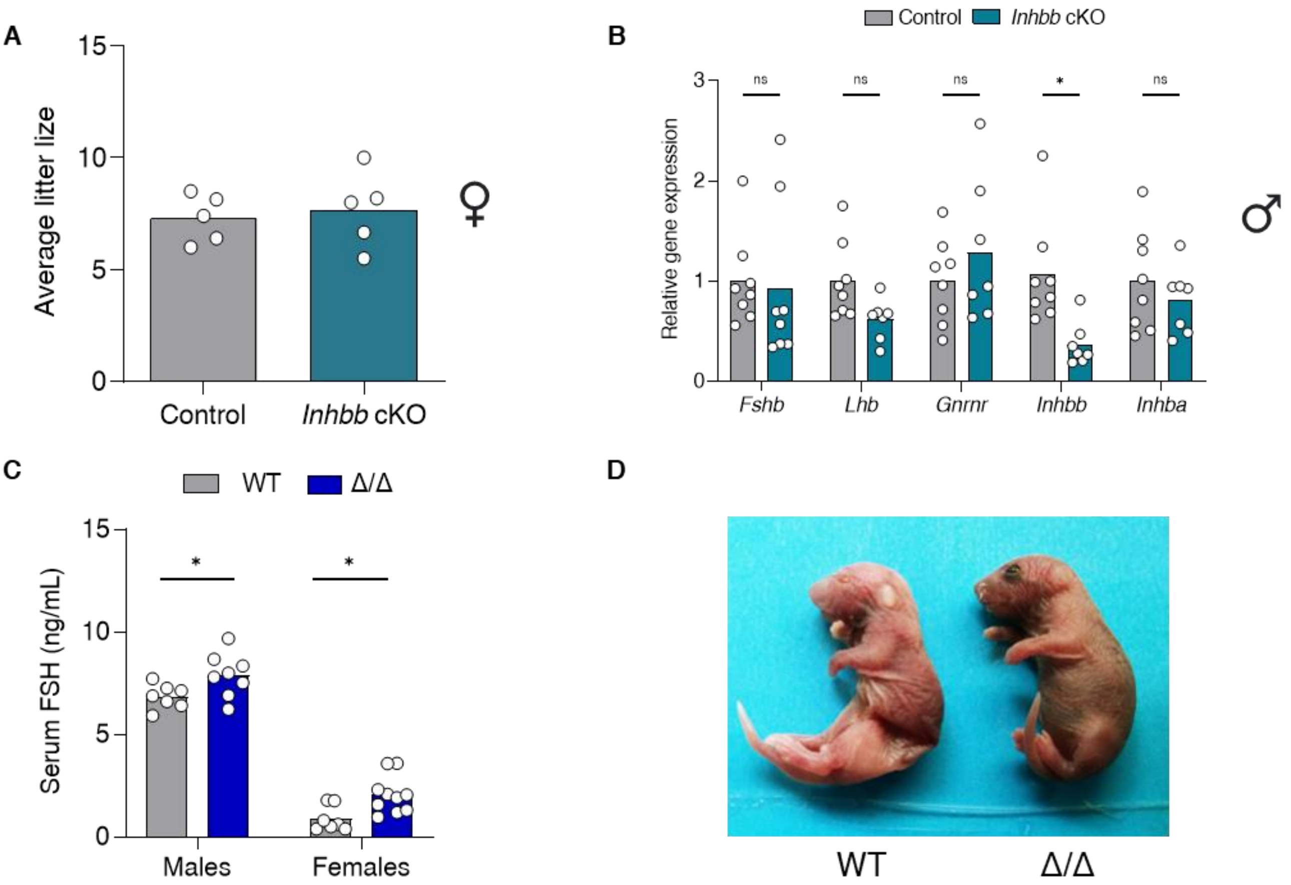


**Fig. S1**: **FSH synthesis is activin B-independent in mice. (A)** Number of pups per litter in 6-month breeding trials of control (gray) and *Inhbb* cKO females (teal). **(B)** Pituitary gene expression in male control or gonadotrope-specific *Inhbb* cKO mice. *Rpl19* was used as a housekeeping gene. **(C)** Serum FSH levels (measured by ELISA) in wild-type (+/+) or *Inhbb* global KO (Δ/Δ) males and females. The *Inhbb*$^{\Delta}$ allele was generated by globally recombining the floxed *Inhbb* allele. Animals were sacrificed at 8 to 10 weeks old. Bar heights are group means. Each circle represents an individual mouse. * *P* < 0.05. ns, non-significant. **(D)** Representative image of eyelid malformations in *Inhbb* KO (Δ/Δ) pups (right).

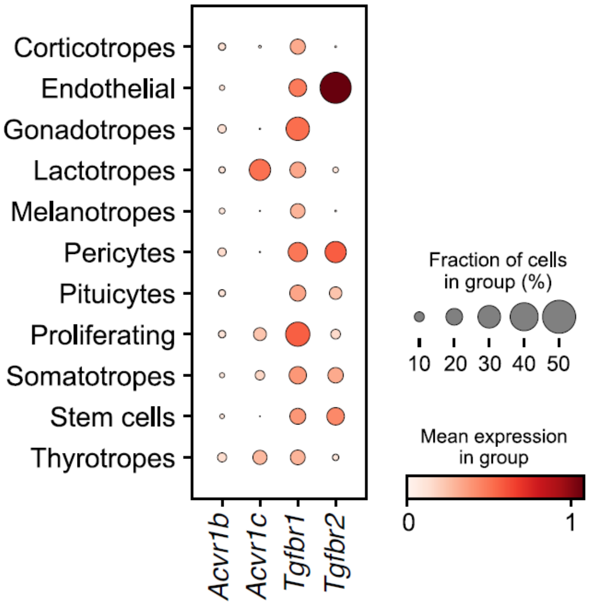


**Fig. S2: ACVR1B and TGFBR1 are expressed in all cell types of the pituitary gland:** Dot plots of *Acvr1b*, *Acvr1c*, *Tgfbr1*, and *Tgfbr2* expression in different cell lineages from snRNAseq of adult female and male murine pituitaries.

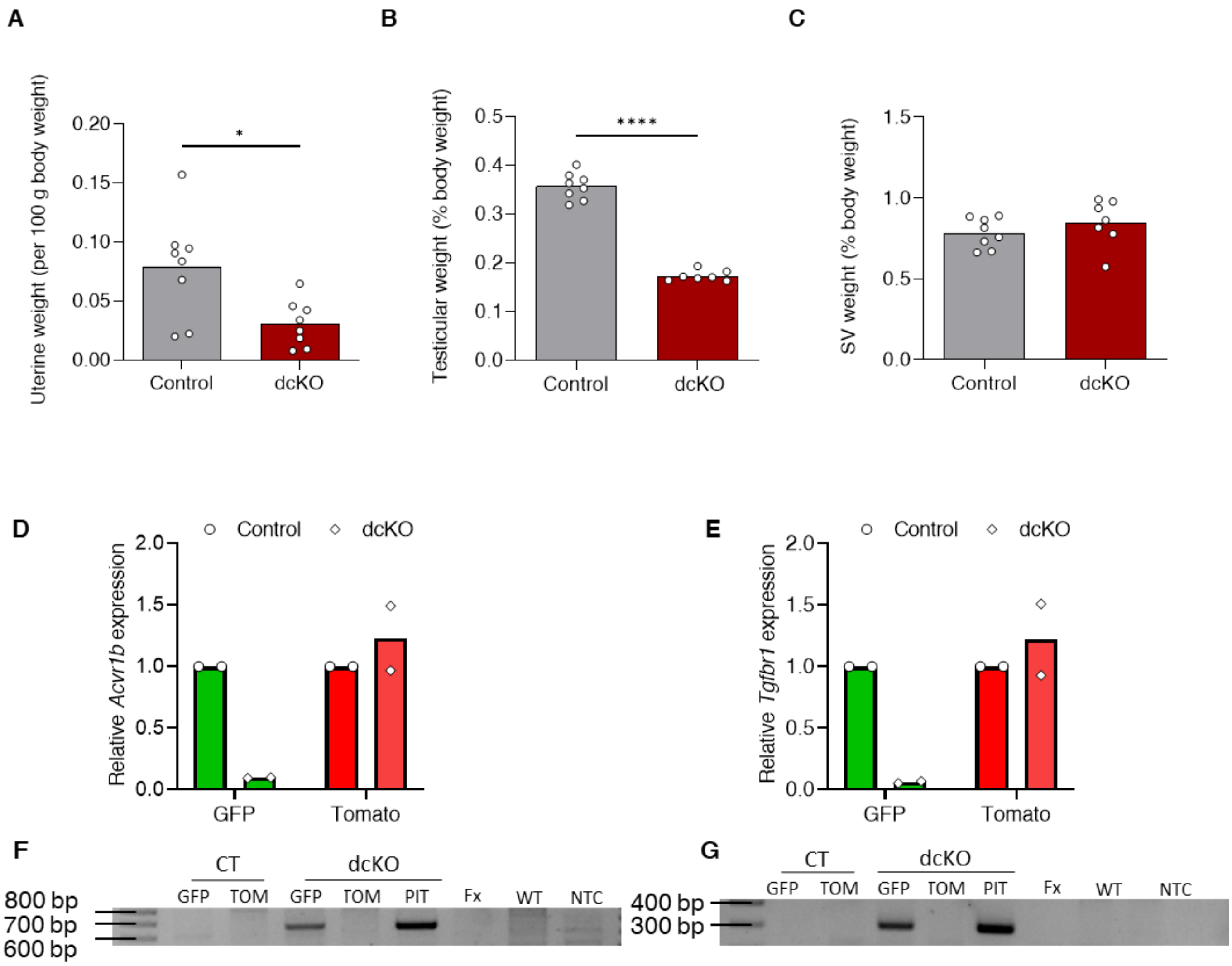


**Fig. S3**: **Characterization of *Acvr1b/Tgfbr1* double cKO mice. (A)** Uterine, **(B)** testicular, and **(C)** seminal vesicle weights (normalized to body weight) in control and gonadotrope-specific *Acvr1b/Tgfbr1* dcKO mice. Bar heights are group means. Each circle represents an individual mouse. * $P < 0.05$, **** $P < 0.0001$. Relative gene expression of **(D)** *Acvr1b* (left) and **(E)** *Tgfbr1* (right) in purified gonadotrope (GFP) versus non-gonadotrope (tomato or TOM) cells from control (CT; *Acvr1b*$^{+/+}$;*Tgfbr1*$^{+/+}$;*Rosa26*$^{mTmG/+}$;*Gnrhr*$^{GRIC/+}$) and dcKO (*Acvr1b*$^{fx/fx}$;*Tgfbr1*$^{fx/fx}$;*Rosa26*$^{mTmG/+}$;*Gnrhr*$^{GRIC/+}$) mice. *Rpl19* was used as housekeeping gene. Bar heights are group means. Each circle represents an independent experiment of a pool of N=3 animals/genotype per condition. PCR of genomic DNA from purified gonadotropes (GFP) and non-gonadotropes (TOM) showing DNA recombination of floxed **(F)** *Acvr1b* and **(G)** *Tgfbr1* alleles. Pituitary glands from dcKO mice (PIT) and tails from floxed (fx) and (WT) wild-type mice were used to show recombination specificity. CT, control; dcKO, double conditional knockout; NTC, no template control.

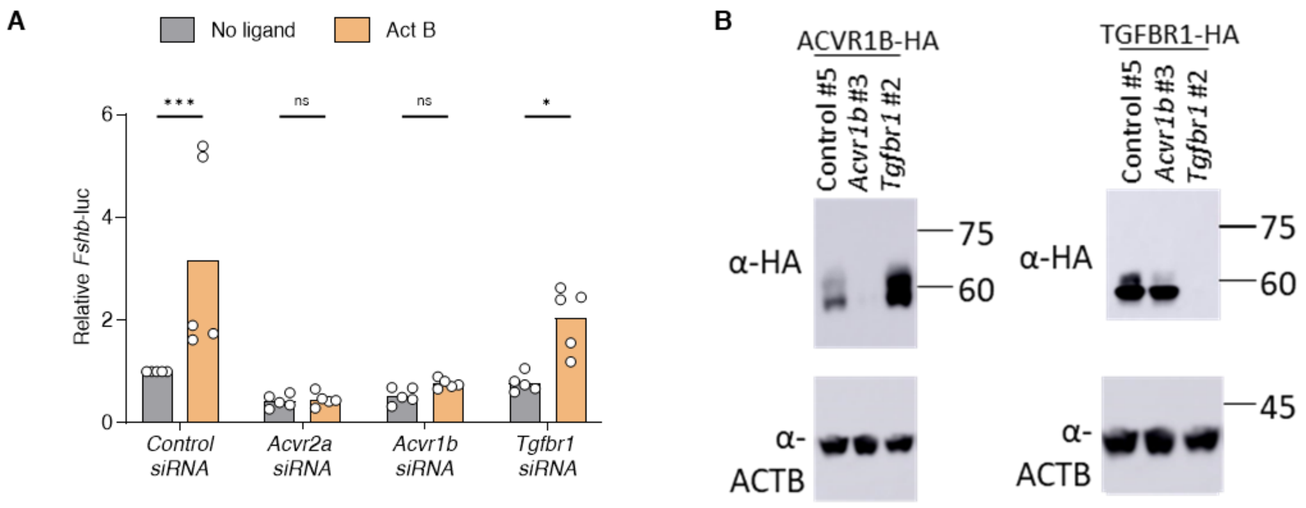


**Fig. S4: Activin B signals via ACVR1B in gonadotrope-like cells. (A)** LβT2 cells were transfected with the murine *Fshb*-luc reporter and 5 nM of control, *Acvr2a*, *Acvr1b*, or *Tgfbr1* siRNA. Cells were treated with media or activin B (2 nM). Bar heights are group means. Each circle represents an independent experiment. * $P < 0.05$, *** $P < 0.001$. ns, non-significant. **(B)** HEK293 cells were transfected with ACVR1B-HA (left) or TGFBR1-HA (right) expression vectors along with 20 nM of control, *Acvr1b*, or *Tgfbr1* siRNA. Protein lysates were immunoblotted against HA. Beta-actin was used as a loading control.

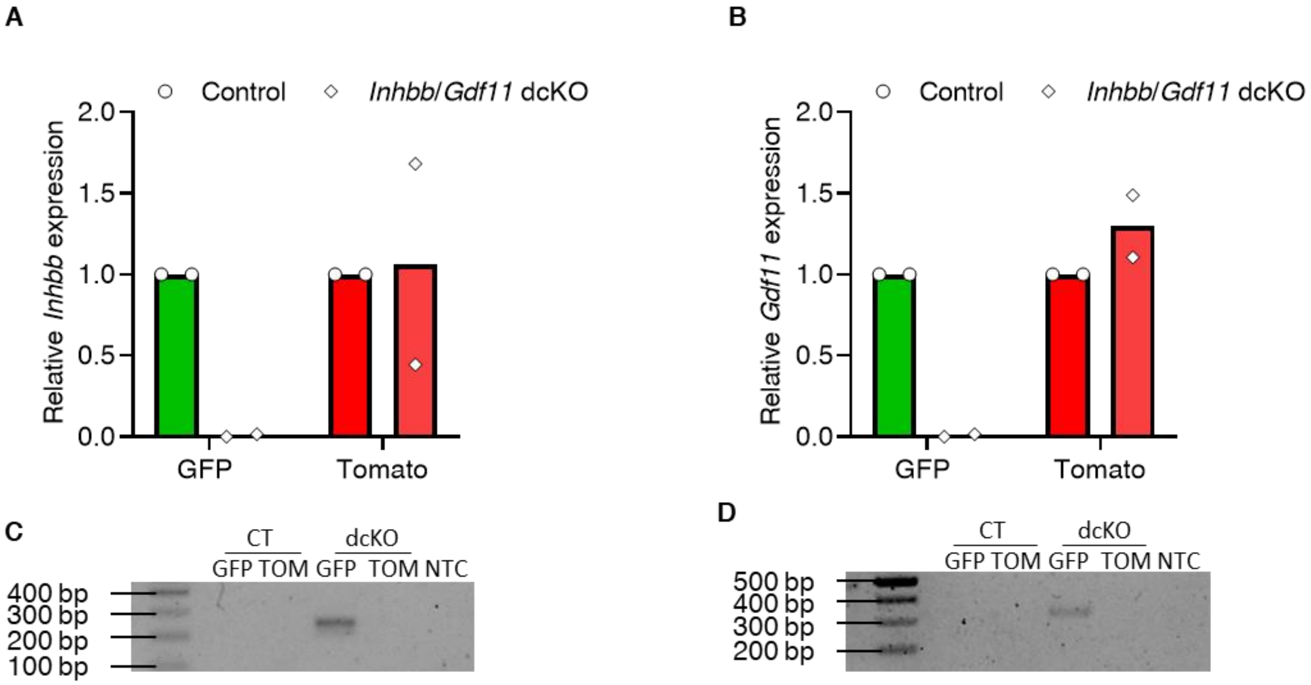


**Fig. S5: Depletion of *Inhbb* and *Gdf11* in gonadotropes of *Inhbb/Gdf11* dcKO mice. (A)** *Inhbb* and **(B)** *Gdf11* expression in purified gonadotrope (GFP) versus non-gonadotrope (TOM) cells from control (*Inhbb*$^{+/+}$;*Gdf11*$^{+/+}$;*Rosa26*$^{mTmG/+}$;*Gnrhr*$^{GRIC/+}$) and *Inhbb/Gdf11* dcKO (*Inhbb*$^{fx/fx}$; *Gdf11*$^{fx/fx}$;*Rosa26*$^{mTmG/+}$;*Gnrhr*$^{GRIC/+}$) mice. *Rpl19* was used as housekeeping gene. Bar heights are group means. Each circle represents an independent experiment of a pool of N=3 animals/genotype per condition. PCR of genomic DNA from purified gonadotropes (GFP) and non-gonadotropes (TOM) showing DNA recombination of floxed **(C)** *Inhbb* and **(D)** *Gdf11* alleles in dcKO cells. CT, control; dcKO, double conditional knockout; NTC, no template control.

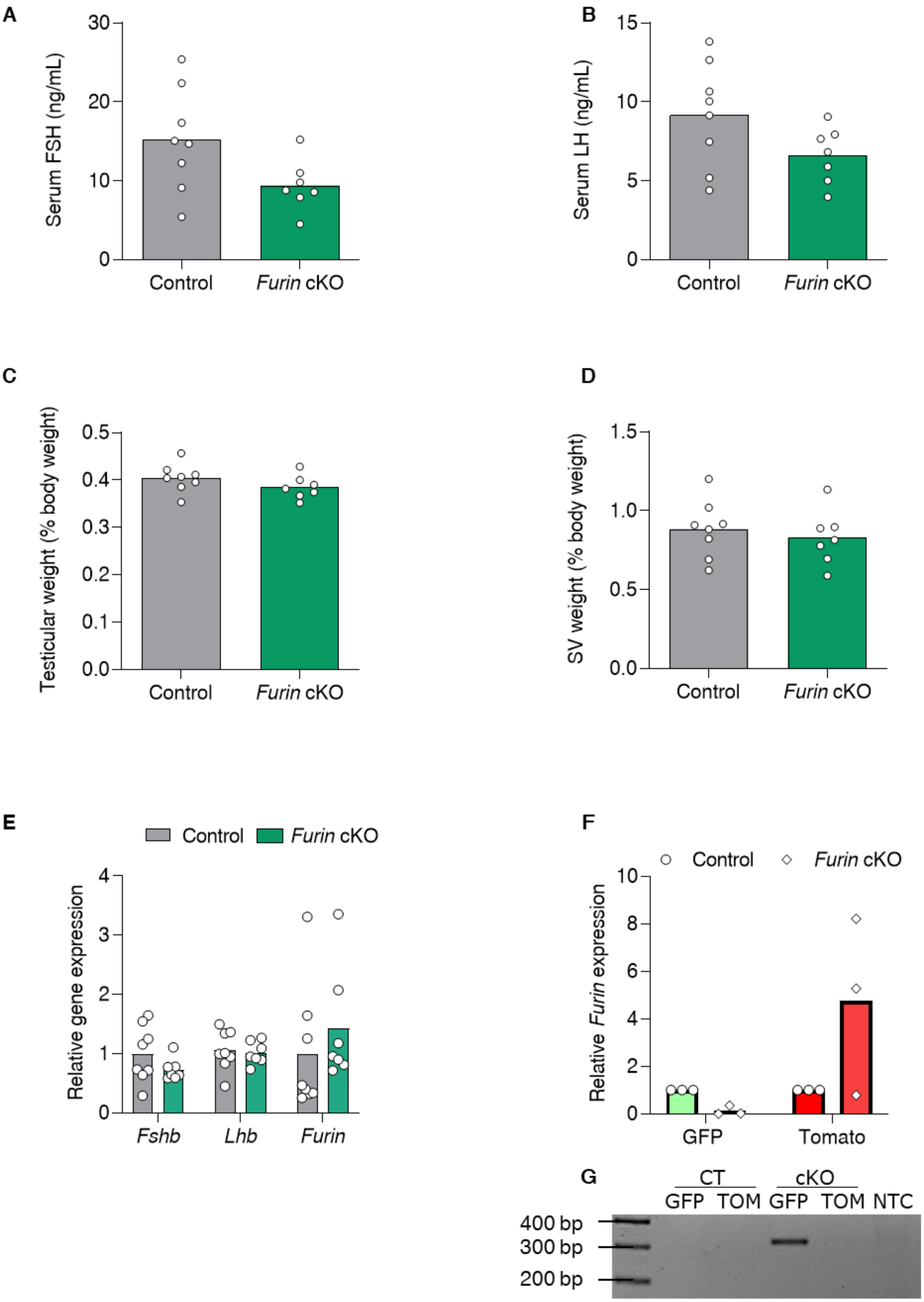
A
Serum FSH (ng/mL)
0
10
20
30
Control
Furin cKO
B
Serum LH (ng/mL)
0
5
10
15
Control
Furin cKO
C
Testicular weight (% body weight)
0.0
0.1
0.2
0.3
0.4
0.5
Control
Furin cKO
D
SV weight (% body weight)
0.0
0.5
1.0
1.5
Control
Furin cKO
E
Control
Furin cKO
Relative gene expression
0
1
2
3
4
Fshb
Lhb
Furin
F
Control
Furin cKO
Relative Furin expression
0
2
4
6
8
10
GFP
Tomato
G
CT
cKO
GFP TOM GFP TOM NTC
400 bp
300 bp
200 bp

**Fig. S6: *Furin* expression in gonadotropes is not necessary for FSH production in male mice.** Serum **(A)** FSH and **(B)** LH levels (measured by ELISA) in control (gray) or gonadotrope-specific *Furin* cKO (green) male mice. **(C)** Testicular and **(D)** seminal vesicle weights (normalized to body weight) in control and *Furin* cKO mice. **(E)** Pituitary gene expression in control and *Furin* cKO mice. *Rpl19* was used as a housekeeping gene. Animals were sacrificed at 8-9 weeks old. Bar heights are group means. Each circle represents an individual mouse. **(F)** Relative *Furin* expression in purified gonadotrope (GFP) versus non-gonadotrope (TOM) cells from control ($Furin^{+/+}$;$Rosa26^{mTmG/+}$;$Gnrhr^{GRIC/+}$) and cKO ($Furin^{fx/fx}$;$Rosa26^{mTmG/+}$;$Gnrhr^{GRIC/+}$) mice. *Rpl19* was used as housekeeping gene. Bar heights are group means. Each circle represents an independent experiment of a pool of N=3 animals/genotype per condition. **(G)** PCR of genomic DNA from purified gonadotropes (GFP) and non-gonadotropes (TOM) showing DNA recombination of floxed *Furin* alleles in cKO gonadotropes. CT, control; cKO, conditional knockout; NTC, no template control

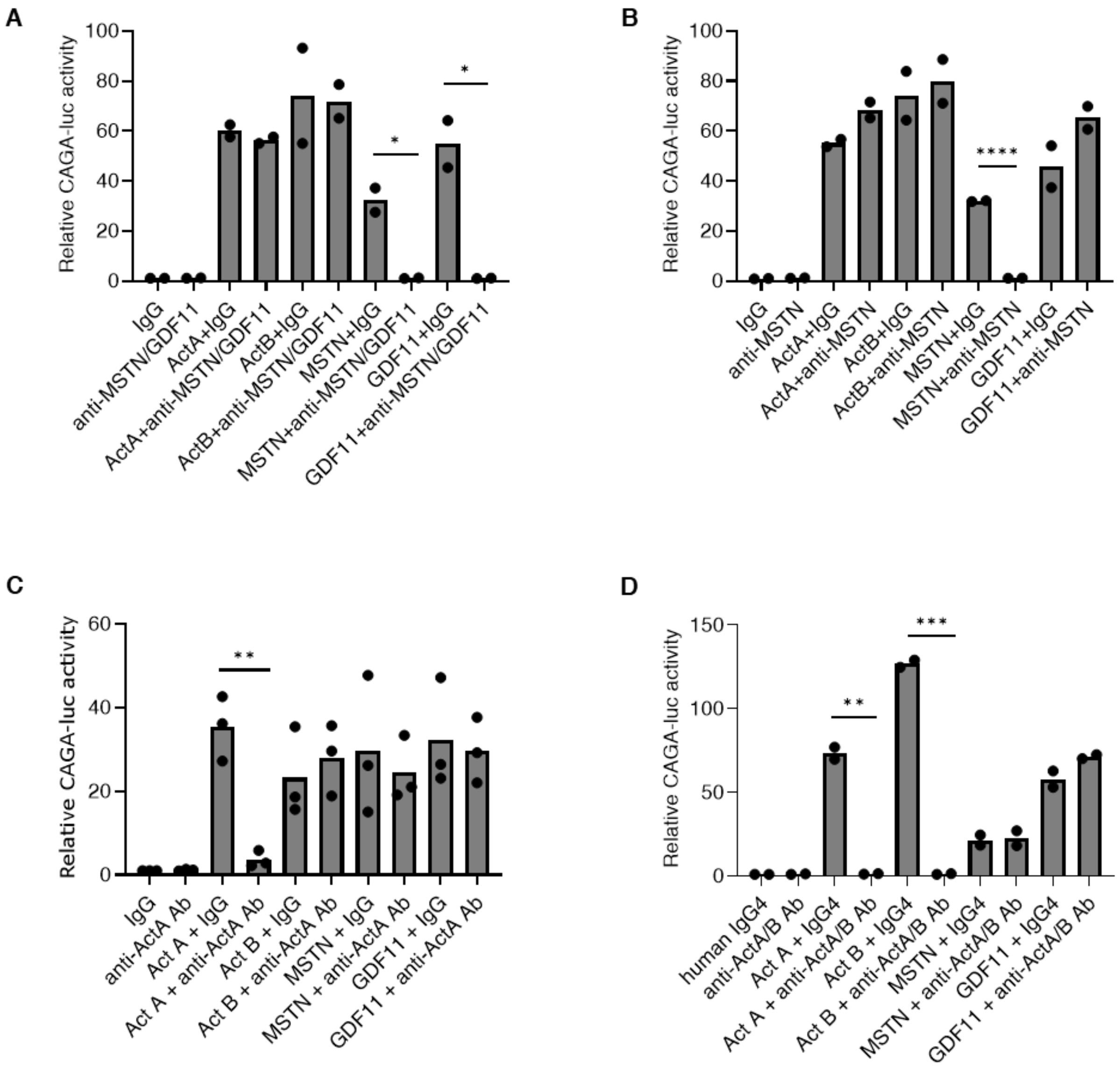


**Fig. S7: Specificity of the neutralizing antibodies.** HEK293-$CAGA_{12}$-luciferase reporter cells were treated with 0.1 nM activin A, activin B, MSTN, or GDF11 $\pm$ 100X $EC_{50}$ of the **(A)** anti-MSTN (RK22) or the **(B)** anti-MSTN/GDF11 antibody (RK35). In **(C)**, cells were treated with 1 nM activin A, activin B, MSTN or GDF11 $\pm$ 100X $EC_{50}$ of ant-ActA (SW101). In **(D)**, cells were treated with 0.1 nM activin A, activin B, MSTN, or GDF11 $\pm$ dual activin A/activin B antibody (REGN16430, 1 mg/mL). Bar heights are group means. Each circle represents an independent experiment. * $P < 0.05$, ** $P < 0.01$, *** $P < 0.001$, **** $P < 0.0001$.

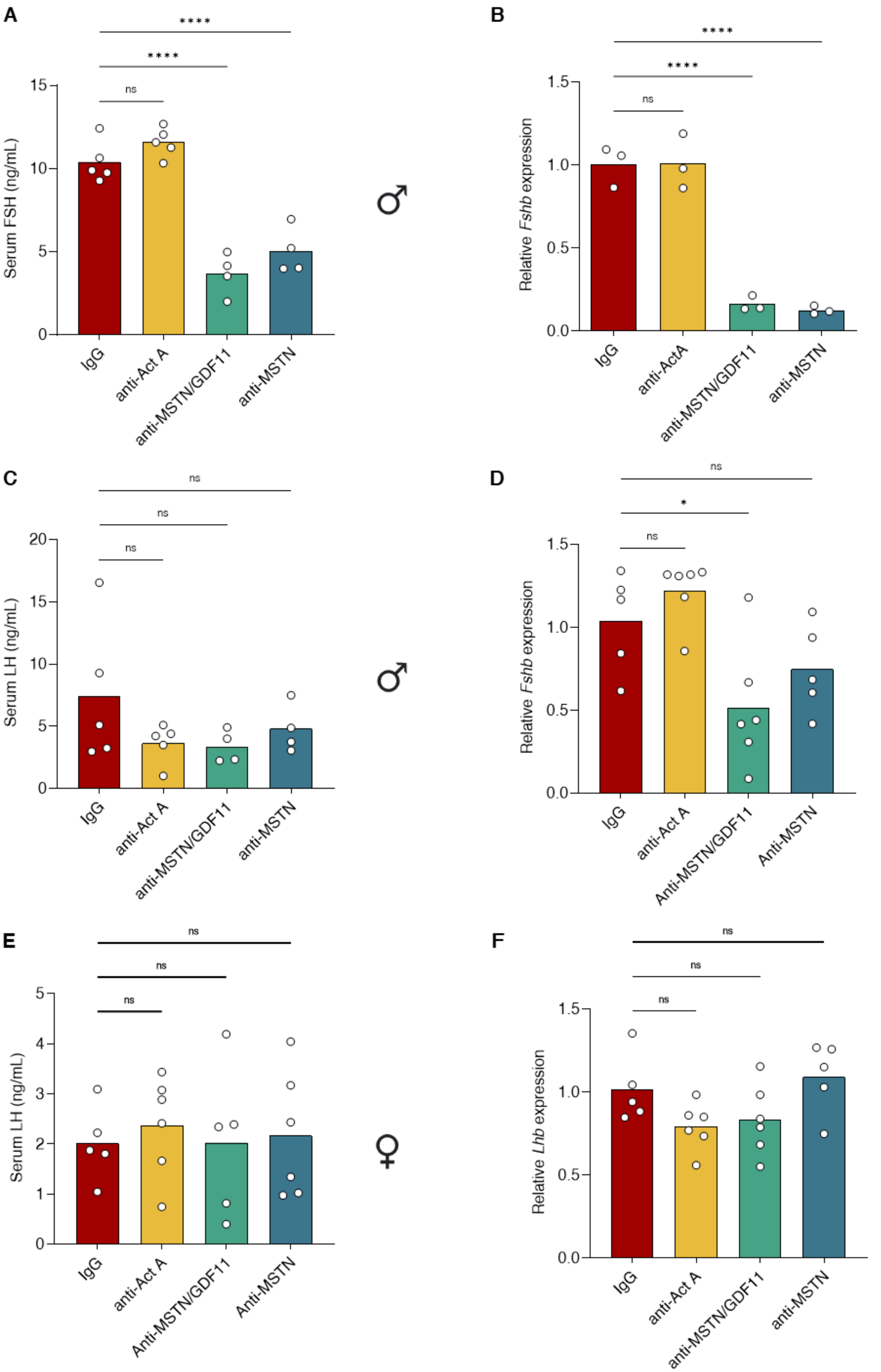
A
B
C
D
E
F
Serum FSH (ng/mL)
Relative Fshb expression
Serum LH (ng/mL)
Relative Lhb expression
IgG
anti-Act A
anti-ActA
anti-MSTN/GDF11
Anti-MSTN/GDF11
anti-MSTN
Anti-MSTN
****
ns
*
♂
♀

**Fig. S8: Myostatin neutralization decreases FSH levels in mice. (A)** Serum FSH levels (ELISA), **(B)** pituitary *Fshb* expression, and **(C)** serum LH levels in male mice 7 days after injections with the indicated neutralizing antibodies. **(D)** Pituitary *Fshb* expression, **(E)** serum LH levels, and **(F)** pituitary *Lhb* expression of female mice at natural ovulation following a single injection i.v. with the indicated neutralizing antibodies. Injected females were paired with wild-type males 7 days post-injection and samples collected on the morning of vaginal plugging. Bar heights are group means. Each circle represents an individual mouse. * $P < 0.05$, **** $P < 0.0001$. ns, non-significant.

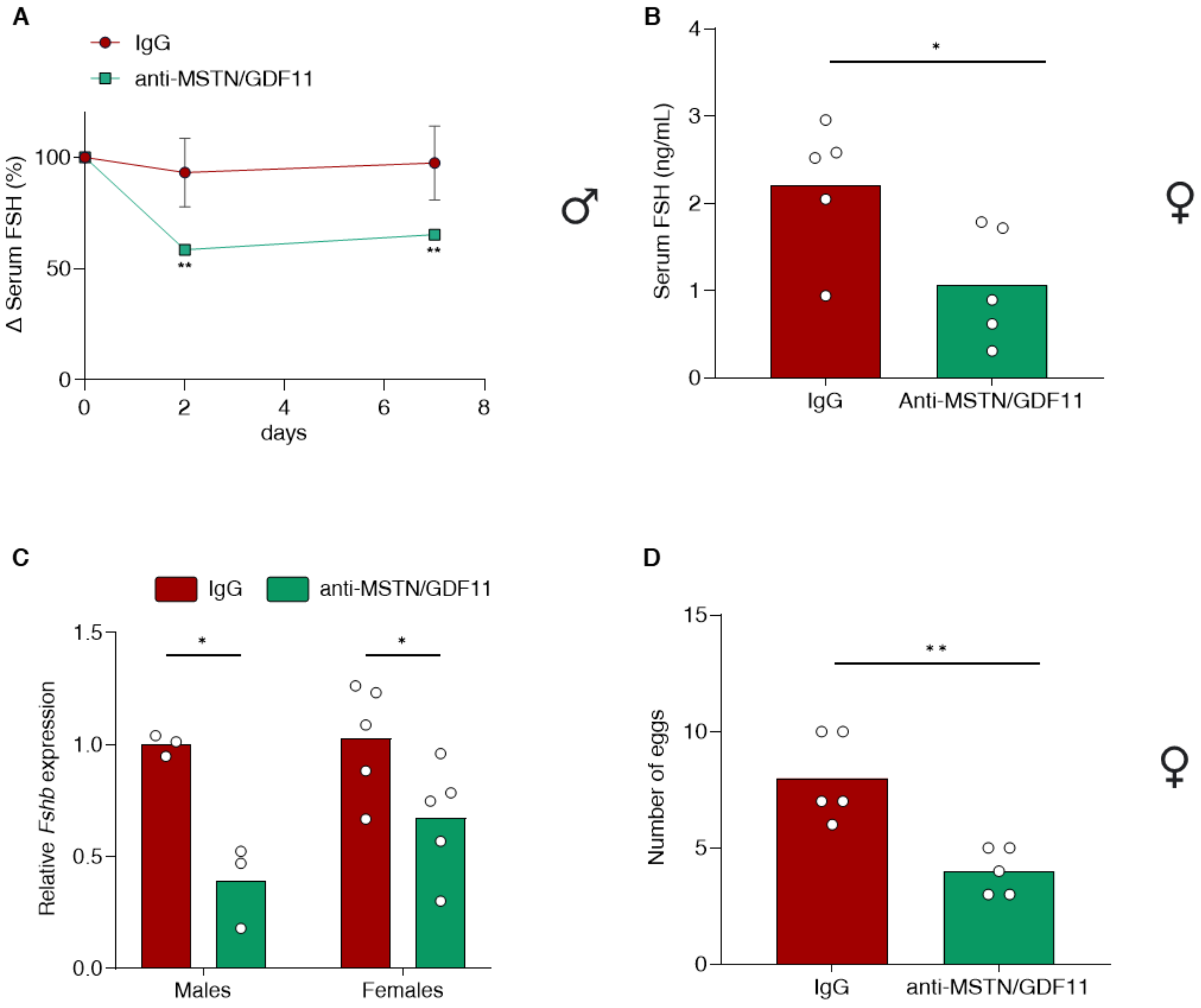


**Fig. S9: GDF11 promotes FSH synthesis in *Mstn* KO mice. (A)** Percentage change in serum FSH levels 2 or 7 days following a single i.v. injection of mouse IgG or anti-MSTN/GDF11 antibody (10 mg/kg) in male *Mstn* KOs. Data were normalized to FSH levels before injection for each mouse. Data represent mean ± SEM (N=3/group). **(B)** Serum FSH levels (ELISA) in females, **(C)** pituitary *Fshb* expression in both sexes, and **(D)** number of eggs ovulated by *Mstn* KO mice at natural ovulation following a single i.v. injection of mouse IgG or anti-MSTN/GDF11 antibody (10 mg/kg). Injected females were paired with wild-type males 7 days post-injection and samples collected on the morning of vaginal plugging. Bar heights are group means. Each circle represents an individual mouse. * $P < 0.05$ and ** $P < 0.01$.

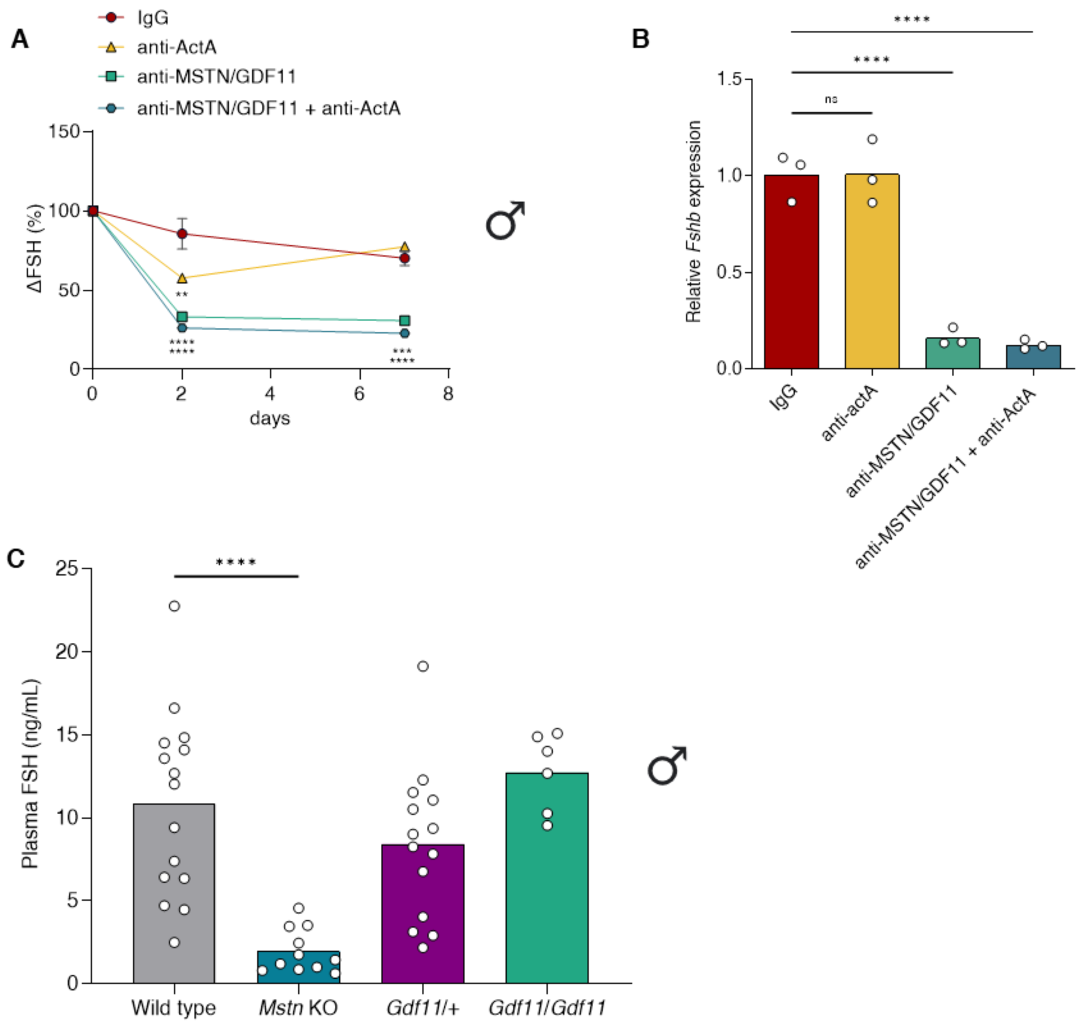


**Fig. S10: Myostatin is a major driver of FSH synthesis and secretion in rodents. A)** Percentage change in serum FSH levels 2 or 7 days and following a single i.v. injection of mouse IgG, anti-activin A, or/and anti-MSTN/GDF11 antibodies (10 mg/kg) in adult male wild-type mice. The total amount of injected antibody was balanced to 20 mg/kg using mouse IgG. Data were normalized to FSH levels before injection for each mouse. Data represent mean ± SEM (N=3/group). **(B)** Pituitary *Fshb* expression in male mice 7 days after injection with the indicated neutralizing antibodies. Bar heights are group means. Each circle represents an individual mouse. **(C)** Plasma FSH levels (ELISA) in male mice in which *Gdf11* was knocked into the *Mstn* locus. Plasma was used from archival material in (*13*). Bar heights are group means. Each circle represents an individual mouse. ** $P < 0.01$, *** $P < 0.001$, **** $P < 0.0001$. ns, non-significant.

**Table S1: Primers for genotyping and DNA recombination.**

| **Table S1** | Reference or sequence |
|---|---|
| *Inhbb* KO | *(24)* |
| *Tgfbr1* genotyping | (*54*) |
| *Tgfbr1* DNA recombination | 5’-AGTCATAGAGCATGTGTTAGAGTC-3’ |
| | 5’-ATTTCTTCTGCTATAATCCTGCAG-3’ |
| *Acvr1b* genotyping/ DNA recombination | (*53*) |
| *Furin* genotyping/ DNA recombination | (*56*) |
| *Gdf11* genotyping/ DNA recombination | (*55*) |
| *Inhbb* genotyping | 5’-AGTCAGGGTCTGGTCAAAGGATCGG-3’ |
| | 5’-GATTGCAGGGAGGATTTGGGGGTAG-3’ |
| *Inhbb* DNA recombination | 5’-AGGTGTGTGCTGTAAGTCCTCACC-3’ |
| | 5’-TGAACTGATGGCGAGCTCAGACC-3’ |
| | 5’-CTTGCCTGTTTGCAGTGGAG-3’ |
| *Mstn* KO | (*38*) |
| Cre-recombinase | (*52*) |
| *Gdf9*-iCre | 5’-TCTGATGAAGTCAGGAAGAACC-3’ |
| | 5’-GAGATGTCCTTCACTCTGATTC -3’ |

**Table S2: Primers for gene expression by qPCR**

| **Table S2** | Reference or sequence |
|---|---|
| *Rpl19*, *Fshb*, *Lhb*, *Cga* and *Gnrhr* | (*25*) |
| *Inhbb*, *Inhba* | (*12*) |
| *Tgfbr1* | 5’-CCTCGAGACAGGCCATTTGT-3’ |
| | 5’-GCCAGCTGACTGCTTTTCTG-3’ |
| *Acvr1b* | 5’-CCCTTGCCGATAATCTCTTG-3’ |
| | 5’-CGCTCCAGGATCTCGTCTAC-3’ |
| *Furin* | 5’-ATCTTCACCAACACCTGGGC-3’ |
| | 5’-ACTGCTCTGTGCCAGAAGTG-3’ |
| *Gdf11* | 5’-AAGAGGACGAGTACCACGCT-3’ |
| | 5’-TTGGGCCTTCAGTACCTTGG-3’ |
